# Lipidomic analysis reveals differences in *Bacteroides* species driven largely by plasmalogens, glycerophosphoinositols and certain sphingolipids

**DOI:** 10.1101/2022.08.11.503646

**Authors:** Eileen Ryan, Belén Gonzales Pastor, Lee A. Gethings, David J. Clarke, Susan A. Joyce

## Abstract

There has been increasing interest in bacterial lipids in recent years due, in part, to their emerging role as molecular signalling molecules. *Bacteroides thetaiotaomicron* is an important member of the mammalian gut microbiota that has been shown to produce sphingolipids (SP) that pass through the gut epithelial barrier to impact host SP metabolism and signal into host inflammation pathways. *B. thetaiotaomicron* also produces a novel family of N-acyl amines (called glycine lipids) that are potent ligands of host Toll-like receptor 2 (TLR2). Here, we specifically examine the lipid signatures of 4 species of gut associated *Bacteroides*. In total we identify 170 different lipids and we report that the range and diversity of *Bacteroides* lipids is species-specific. Multi-variate analysis reveals that the differences in the lipid signatures are largely driven by the presence/absence of plasmalogens, glycerophosphoinositols and certain SP. Moreover, we show that, in *B. thetaiotaomicron*, mutations altering either SP or glycine lipid biosynthesis results in significant changes in the levels of other lipids suggesting the existence of compensatory mechanisms required to maintain the functionality of the bacterial membrane.

**Importance:** *Bacteroides* are important beneficial members of the gut microbiome that produce lipids that can function as cross-kingdom signalling molecules. We describe, for the first time, a comprehensive and qualitative comparison of the lipid signatures of 4 important *Bacteroides* species. We identify a group of *Bacteroides* core lipids and uncover species-specific differences in plasmalogen, glycerophospholipid and SP metabolism with more subtle differences observed in glycine lipid production. This data will provide a useful platform for the further characterisation of the lipid-based host-microbe dialogue and the influence of microbial lipids on host health and disease states.

## Introduction

Bacterial lipids have recently emerged as influential contributors to the microbe-host molecular dialogue (Chandler & Ernst, 2017; Wei *et al*., 2019; Lamichhane *et al*., 2021). Lipids are hydrophobic or amphipathic small molecules found in all living cells, including bacteria, with important functions in membrane structure, energy storage and cell signalling (Ridgeway & McLeod, 2015). Based on their chemical structures and biosynthetic origins, lipids have been grouped into eight categories; fatty acyls (FA), glycerolipids (GL), glycerophospholipids (GP), sphingolipids (SP), sterol lipids (ST), prenol lipids (PR), saccharolipids (SR), and polyketides (PK) and, within each category, there are distinct classes and subclasses (Fahy *et al*., 2005; Liebisch *et al*., 2020). To date, over 40,000 biologically relevant lipids (mostly mammalian) are listed to the LIPID MAPS Structural Database (Sud *et al*., 2007). However, there is growing interest in cataloguing non-mammalian lipids, including those produced by gut microbes, prompted, in part, by the identification of some bacterial lipids as molecular signals (Chandler & Ernst, 2017; Lamichhane *et al*., 2021; Hou *et al*., 2022).

*Bacteroides* are early colonisers of the mammalian gut, establishing stable, long-term, and generally beneficial interactions with their human host (Wexler & Goodman, 2017). *Bacteroides* have been shown to produce a variety of important bioactive lipids including sphingolipids (SP) and N-acyl amines called glycine lipids (Wieland Brown *et al*., 2013; An *et al*., 2011; An *et al*., 2014; Brown *et al*., 2019; Clark *et al*., 2013; Wang *et al*., 2015; Cohen *et al*., 2017; Nemati *et al*., 2017; Nichols *et al*., 2020a, 2020b). It is now well recognised that *Bacteroides* are one of only a few bacterial genera that produce SP (Heaver *et al*., 2018, Brown *et al*., 2019, Johnson *et al*., 2020). *B. fragilis* generates a bioactive SP (α-galactosylceramide, α-GalCer) which binds to the antigen-presenting protein, CD1d, thus influencing the number and function of natural killer T cells (NKT-cells) in the intestine, with consequences in the progression of a murine model of colitis (Wieland Brown *et al*., 2013; An *et al*., 2014). In another study, the lack of bacterial SP production was shown to promote intestinal inflammation with concurrent changes in the host SP pool (Brown *et al*., 2019). Indeed, SP produced by *Bacteroides* have been shown to cross the epithelial barrier and impact hepatic SP pools (Johnson *et al*., 2020).

Glycine lipids are a family of lipids derived from the initial *N*-acylation of glycine that results in the production of a mono-acylated glycine molecule called commendamide (Cohen *et al*., 2017; Lynch *et al*., 2017). Commendamide is further modified by an *O-*acylation resulting in a diacylated glycine (GlyL) with additional modifications resulting in the generation of a family of glycine lipids that includes a serine-glycine dipeptido-lipid (flavolipin, FL) and a large complex glycine lipid called Lipid 1256 (Bill *et al*., 2021; Nichols *et al*., 2021; Nichols *et al*., 2020a, 2020b; Lynch *et al*., 2019). Commendamide was originally identified in a screen to identify agonists of GPCR G2A/132 that result in increased levels of NF-kB expression (Cohen *et al*., 2017). GlyL, FL and Lipid 1256 have been reported to signal to eukaryotic cells by engaging TLR2, promoting the production of pro-inflammatory cytokines (Clark *et al*., 2013; Wang *et al*., 2015; Nemati *et al*., 2017; Nichols *et al*., 2020a, 2020b).

It is clear, therefore, that *Bacteroides* produce unique lipids with the potential to signal to the mammalian host. However, a comprehensive examination of the *Bacteroides* lipid signature has not been carried out and this limits a full appreciation of their potential in the host-microbe dialogue. In the present study we describe and compare the lipid signatures of 4 important species of *Bacteroides*: namely *B. thetaiotaomicron, B. fragilis, B. ovatus* and B. *vulgatus* (recently elevated to *Phocaeicola vulgatus*), with a particular focus on the bioactive SP and glycine lipids. We identify 170 different lipids and show that the lipid signatures vary in a species-dependent manner. In addition, we show that mutations in SP or glycine lipid biosynthesis significantly changes the lipid signature of *B. thetaiotaomicron* and these compensatory changes need to be considered when studying the role of these lipids in *Bacteroides* and in the microbe-host dialogue.

## Materials & Methods

### Materials

Organic solvents (Supelco 2-propanol and acetonitrile) used for the extractions/precipitations and mobile phase preparation were hypergrade for LC-MS LiChroslv® and obtained from Merck (Darmstadt, Germany). Buffers used for mobile phase preparation were from Fisher Chemical Optima™ LC-MS grade Formic Acid from Fisher (Leicestershire, UK) and LC-MS grade, LiChropur™ ammonium formate from Merck (Darmstadt, Germany). Internal standard was N-palmitoyl-D-erythro-sphingosylphosphorylcholine (16:0 SM, Avanti Polar Lipids, powder) was purchased from Merck (Darmstadt, Germany).

### Bacterial strains and growth conditions

*B. thetaiotaomicron* VPI-5482 Δtdk (gift from Eric Martens), *B. fragilis* 638R (gift from Eduardo Rocha), *B. ovatus* ATCC8483 Δtdk (gift from Eric Martens and Nicole Koropatkin) and *B. vulgatus* ATCC8482 (gift from Eric Martens) were anaerobically cultured at 37°C in brain heart infusion (BHI) medium (Sigma) supplemented with hemin (5 μg ml^-1^), 0.1% (wt/vol) cysteine, and 0.2% (wt/vol) sodium bicarbonate. The *B. thetaiotaomicron* Δ*SPT* mutant was a gift from Eric Brown (Brown *et al*., 2019), whilst the *B. thetaiotaomicron* Δ*gls*B mutant was constructed as previously described (Lynch *et al*., 2019).

### Lipid Extraction

Bacteria were inoculated in BHIS browth and, after 24h, cultures were normalised to 1 OD_600_, and pelleted by centrifugation at 8,000 rpm x 10 minutes. Pellets were washed twice with phosphate bufferered saline (PBS). Bacterial pellets were subject to a single phase isopropanol lipid extraction and protein percipitation procedure as descirbed by Sarafian *et al*. (2014). Briefly, washed pellets were resuspended in isopropanol (to a final density of 1 OD_600_ ml^-1^) with added internal standard (16:0 SM, 1µg/mL) and vortexed for 30 sec. Samples were incubated at room temperature for 10 min, with occassional mixing, and stored overnight at - 20°C. The supernatent was collected following centrifugation for 20 minutes at 14,000 g and stored at -80°C for LC-MS analysis

### LC-MS Conditions

Bacterial isopropanol lipid extracts were analysed using Waters Xevo™ G2-XS QTOF Mass Spectrometer coupled to a Waters ACQUITY™ UPLC™ system. Extracted samples were injected (5 μL injection volume) onto a ACQUITY CSH™ column (100 mm × 2.1 mm, 1.7 μm; Waters) at 55 °C with a flow rate of 400 μL/min. The mobile phases consisted of phase A (acetonitrile/water (60:40, v:v) with 10 mM ammonium formate and 0.1% formic acid) and it was subject to gradient with mobile phase B (isopropanol/acetonitrile (90:10, v:v) mixed with 10 mM ammonium formate and 0.1% formic acid) (Table S1 in the Supporting Information details the gradient parameters).

Mass spectrometry was performed under both positive and negative ESI modes using the following paramaters: Acquistion mode: MS^E^; acquistion range: from m/z 100 to 2000 acquisition time: 1 sec/scan; source temperature: 120°C; desolvation temperature: 550°C; nitrogen gas flow: 900 L/h; capillary voltage: 2.0 kV (positive mode) or 1.5 kV (negative mode); cone voltage: 30V. For both ionization modes, leucine enkephalin (m/z 556.2771 in ESI+, m/z 554.2615 in ESI-) was continuously infused at 30 μL/min and sampled every 30 seconds for lock mass correction.

### Characterisation of Plasmalogens

Acid hydrolysis was applied to confirm the presence of plasmalogens in *B. thetaiotamicron* extracts according to Murphy *et al*. (1993). Briefly, isopropanol was removed from lipid extracts under nitrogen, eppendorfs were inverted over five drops of concentrated HCl in a test tube cap for 5 minutes. This caused the complete hydrolysis of the vinyl ether bond of the plasmalogens while the diacyl ester bonds remain intact. The samples were re-extracted with isopropnal then reanalysed by LC-MS as described above.

### MS data processing & statistical analyses

For targeted analysis, MS data was processed using the open-source, Skyline-daily (beta) freeware (MacCoss Lab Software, USA). An in-house *Bacteroides* lipid library was constructed from mining the existing literature, to include classes of: sphingolipids, *N*-acyl amines, fatty acids, and glycerophospholipids with cross checking of raw data in both ionisation modes for accuarte mass (where delta mass < 5 ppm), retention time, and MS/MS fragmentation where possible. This database, together with LC-MS raw data files were uploaded into Skyline and processed following lock mass correction. Both positive and negative mode data were processed, following application of a correction factor (based on peak intensity of several common lipids detected in both modes) for both negative mode data and positive mode data, then normalised to the internal standard. Data was log transformed using MetaboAnalyst 5.0 wed-based platform (Xia *et al*., 2009; Pang *et al*., 2021), prior to multivariate (i.e. PCA, heatplots, ANOVA) and univariate (volcano plots) analysis. Customised python scripts were used to produce volcano plots and GraphPad Prism 5.0 to produce bar-charts and for regression analysis. For untargeted analysis, raw LC-MS data files werefirst processed (peak alignment) using Progenesis QI (Waters Corporation, UK) and then imported into MetaboAnalyst 5.0 for median normalisation, log transformation and multivariate analyses for both positive mode and negative mode data.

## Results

### The lipid signatures of Bacteroides are species-specific

Mass spectrometry applications and analyses reveals that a diverse range of lipids are associated with gut resident representatives of *Bacteroides* (see Table S2 for a full list of the lipids identified in this study). Multivariate analysis indicates that *B. vulgatus* and *B. fragilis* lipid signatures co-cluster, whilst *B. thetaiotaomicron* and *B. ovatus* lipid signatures occupy distinct positions (Figure 1A & B). This analysis suggests that *B. vulgatus* and *B. fragilis* may have similar lipidomes that are distinct from both *B. thetaiotaomicron* and *B. ovatus*.

**Figure 1:**
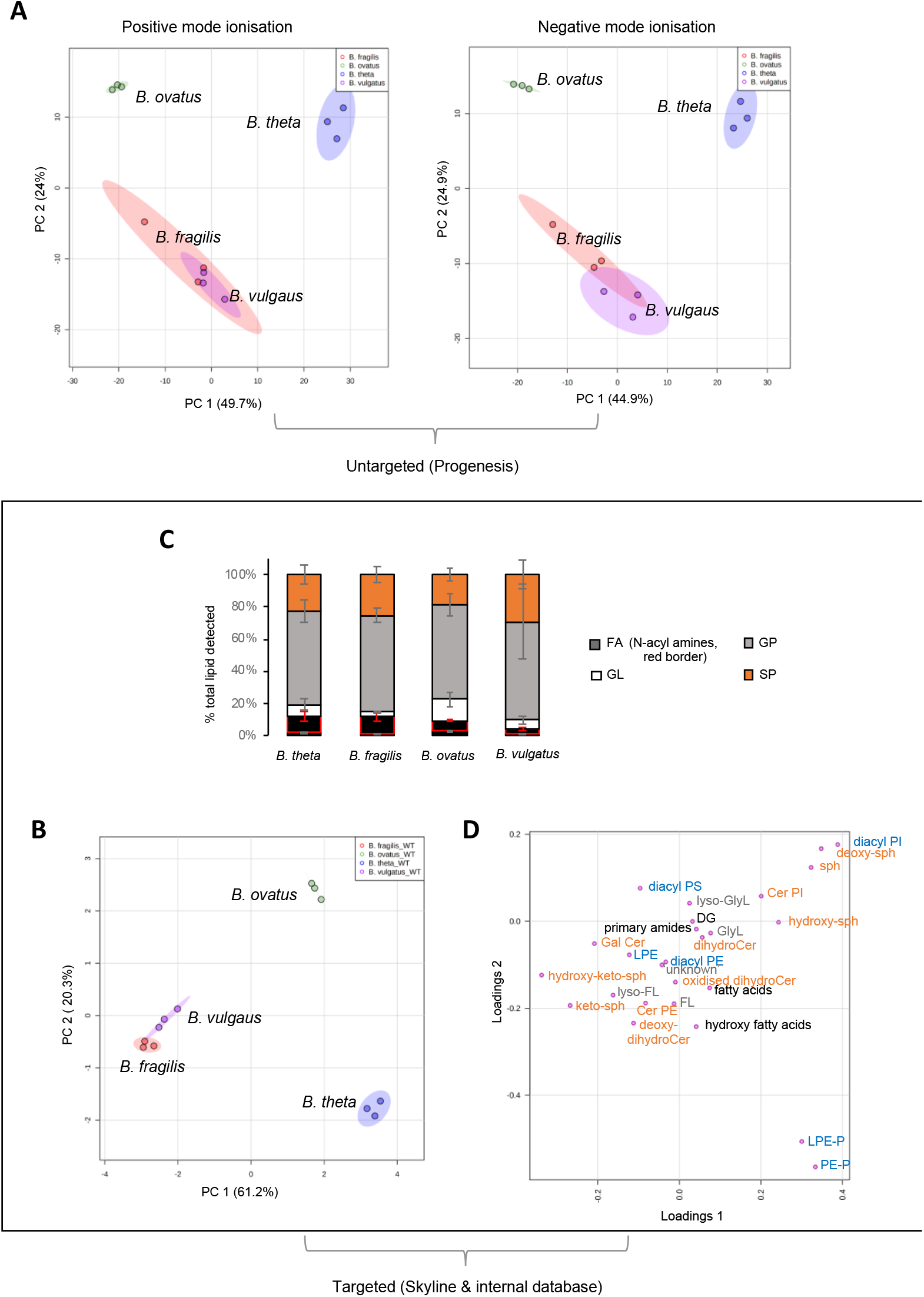
LC-MS and multivariate analysis suggest that *B. fragilis* and *B. vulgatus* may have similar lipidomes that are distinct from both *B. thetaiotaomicron* and *B. ovatus*. (A) PCA scores plot generated from untargeted lipid analysis from either positive or negative mode ionisation data; LC-MS raw data was imported into Progenesis QI, data was median normalised and log transformed using MetaboAnalyst 5.0 wed-based platform prior to multivariate analysis B) PCA scores plot generated from targeted lipid analysis; LC-MS raw data was uploaded into Skyline and processed against an internal lipid database in both positive and negative ionisation modes, following a correction factor applied to negative mode data, it was combined with positive mode data and then normalised to the internal standard, data was log transformed using MetaboAnalyst 5.0 wed-based platform prior to multivariate analysis. (C) Relative distribution of fatty acyls (FA), glycerolipids (GL), glycerophosholipids (GP) and sphingolipids (SP) among the four Bacteroides species. Data are the mean ± SD of three independent biological experiments (D) Loadings plot of B reveals **differences in the lipid profile of Bacteroide wild-type isopropanol extracts are driven largely by plasmalogens (P), diacyl glycerophosphoinositols (PI) and certain SP**. PS: glyerophosphoserines; PE: glycerophosphoethanolamines; LPE: lyso PE; Cer: ceramides; Gal: galactosyl; sph: sphinganines; GlyL: glycine lipids; FL: flavolipins

Targeted LC-MS based lipidomics reveals the diversity of lipids present in *Bacteroides*. In total, we identified 170 individual lipids distributed across 4 lipid categories or 26 lipid ‘subgroups’ (as defined in LIPID MAPS) (Figure 1C). The *Bacteroides* lipidome is dominated by glycerophospholipids (GP) and SP with smaller contributions from fatty acyls (FA) and glycerolipids (GL) (see Figure 1C). Targeted analysis also reveals that the observed differences in the lipid profile between *Bacteroides* species are largely driven by species-dependent signatures related to the presence of plasmalogens (P), diacyl glycerophosphoinositols (PI) and certain SP (Figure 1D). In contrast fatty acids, hydroxy fatty acids, *N*-acyl amines, DG, glycerophosphoethanolamine (both diacyl PE and lyso PE (LPE)), diacyl glycerophosphoserine (PS), dihydroceramide (dihydroCer) and dihydroCer phosphoethanolamines (Cer PE) represent relatively stable core lipids found in all species of *Bacteroides* tested in this study (Figure 1D).

### *N*-acyl amines comprise a signifcant proportion of the fatty acyl (FA) component of *Bacteroides* lipids

The FA category of LIPID MAPS includes fatty acids and *N*-acyl amines. *B. thetaiotaomicron* and *B. fragilis* have the highest representation in the FA category, which inlcude hydroxy fatty acids (C15 to C17), and *N*-acyl amines (Figure 1C). *N*-acyl amines comprising GlyL and FL with varying acyl chain lengths and their respective mono-acylated derivatives (mono-GlyL, mono-FL) were detected in all four *Bacteroides* species (see Figure 2).

**Figure 2:**
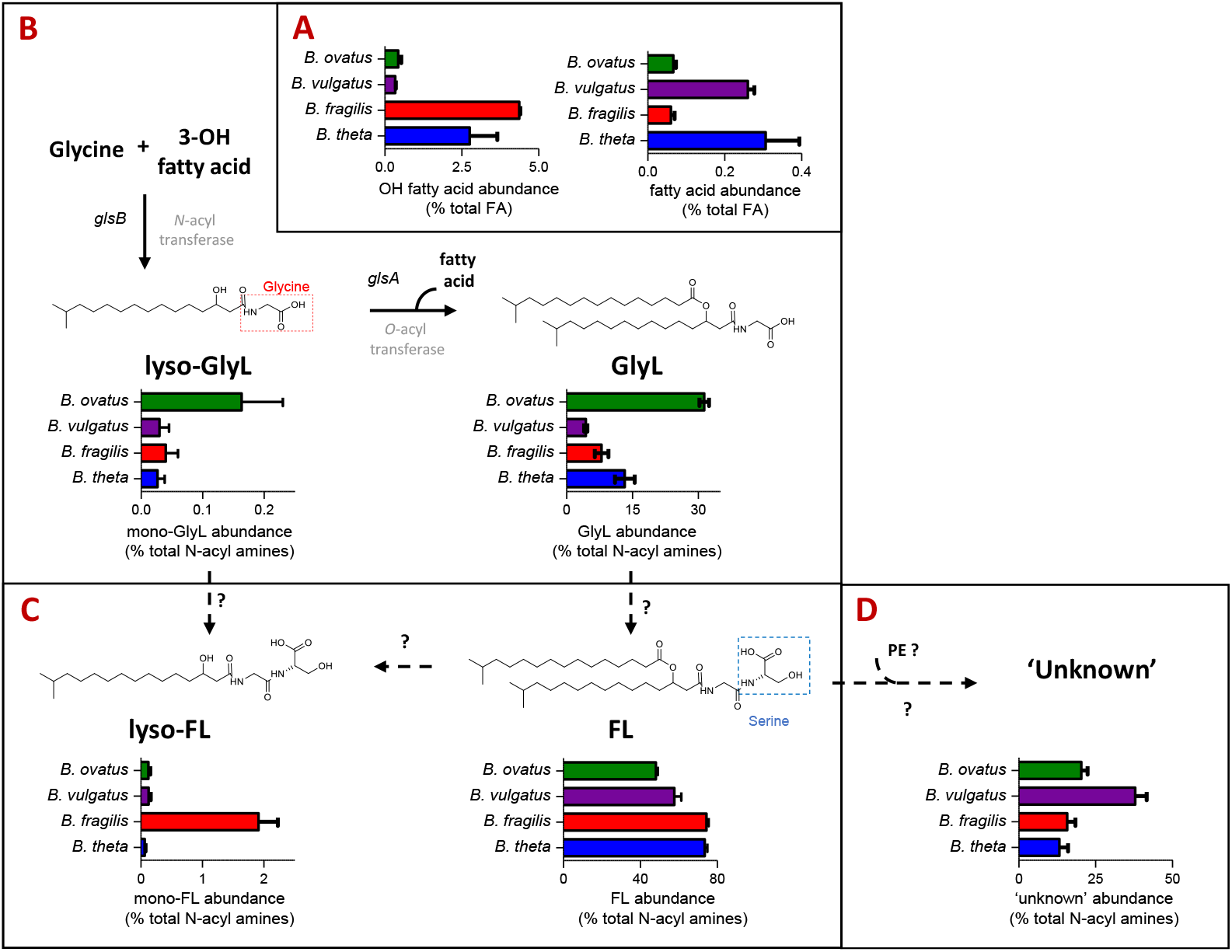
*N*-acyl amines comprise a significant proportion of the fatty acyl (FA) component of *Bacteroides* lipids with some clear quantitative differences observed between species. (A) Relative abundance (expressed as % total FA) of hydroxy fatty acids and fatty acids, among the four *Bacteroides* species tested. Hydroxy fatty acids and fatty acids are the building blocks to more complex *Bacteroides* lipids including *N*-acyl amines shown in B-D. (B) Relative abundance (expressed as % total *N*-acyl amines) of mono-(lyso-GlyL) and diacylated glycine lipids (GlyL) generated from *N*-(encoded by gls*B* gene) and *O*-(encoded by gls*A* gene) acyl transferase (Lynch *et al*., 2019), respectively, in the four Bacteroides species. (C) Relative abundance (expressed as % total *N*-acyl amines) of mono- (lyso-FL) and diacylated flavolipins (FL) in the four Bacteroides species. The exact biosynthesis of FL is unknown but hypothetically they could be synthesised from the respective glycine lipid precursors by attaching serine to the terminal glycine moiety (Nichols *et al*., 2020). (D) Relative abundance (expressed as % total *N*-acyl amines) of the ‘unknown’ lipids predicted to be generated from FL and glycerophosphoethanolamines (PE) in the four Bacteroides species.

GlyL represented between 4% of *B. fragils* and 32% of *B. ovatus* total *N*-acyl amines detected (Figure 2C). For the most part, the major GlyL was the previously reported GlyL at m/z 568 in the positive ion mode, however *B. ovatus* contained similar amounts of a different glycine lipid at m/z 554 in positive ion mode, presumably with a shorter carbon chain length (Figure S1B). For the most part lyso-GlyL (also known as commendamide) was a minor component of the lipid signatures detected for all species (Figure 2C).

In general, FL accounted for most of the *N*-acyl amines detected (Figure 2D) and this lipid was abundantly represented amongst all *Bacteroides* species, with lyso-FL approx. 20 times more abundant in the *B*.*fragilis* lipid signature (Figure 2D). The best characterised FL, at m/z 655 in positive ion mode, is also known as Lipid 654 (FL-654), owing to its molecular weight in the negative ion mode, and this was the most abundant *N*-acyl amine in extracts from *B. fragilis* and *B. vulgatus* whilst the shorter carbon chain length FL, at m/z 641 in the positive ion mode, was the most abundant in *B. thetaiotaomicron* and *B. ovatus* (Figure S1A). These different chain lengths may indicate important strain specific differential signalling.

In addition, a series of ‘unknown’ but predicted *N*-acyl amines with m/z values (in positive ion mode) of 1230.9207 (Unknown_1231), 1244.9363 (Unknown 1245), 1258.9520 (Unknown 1259) and 1272.9676 (Unknown_1273) were also detected. *B. vulgatus* contained a relatively higher proportion of these ‘unknown’ lipids (Figure 2E) with Unknown_1259 and Unknown_1273 representing the most abundant (Figure S1C). Therefore, the profile of *N*-acyl amines is qualitatively similar across all of the examined *Bacteroides* although there are some quantitative differences that, given the important signaling role of the glycine lipid family, may be physiologically important.

### Dihydroceramide phophoethanolamine (Cer PE) is the most abundant sphingolipid (SP) detected in all four *Bacteroides*

In this study, SP were found to represent between 19% *(B. ovatus*) and 29% (*B. vulgatus*) of the total lipids detected (Figure 2A). The SP biosynthetic pathway in *Bacteroides* has been recently described and dihydroCer has been identified as a key intermediate (Stankeviciute *et al*., 2022; Lee *et al*., 2022). In our study dihydroCer was shown to account for approx. 20% of the SP fraction in *B. ovatus, B. vulgatus* and *B. thetaiotaomicron* (Figure 3C). However, in *B. fragilis*, dihydroCer accounted for only 2% of the SP fraction (Figure 3C). Instead, *B. fragilis* appears to accumulate higher levels of keto-sphinganines (keto-sph) which are upstream intermediates in the SP biosynthetic pathway (Figure 3A). Moreover there were detactable levels of sphinganine (sph) and deoxy-sph in both *B. thetaiotamicron* and *B. ovatus* (Figure 3A). These intermediates may be generated by the reduction of keto-sph by a keto-reductase and/or the hydrolysis of dihydroCer by ceramidases (Figure 3A & C). Recent studies have suggested that there may be multiple pathways for SP biosynthesis in *Bacteroides* and the varying levels of intermediates detected across the 4 species examined in this study does support this observation (Stankeviciute *et al*., 2022; Lee *et al*., 2022).

**Figure 3:**
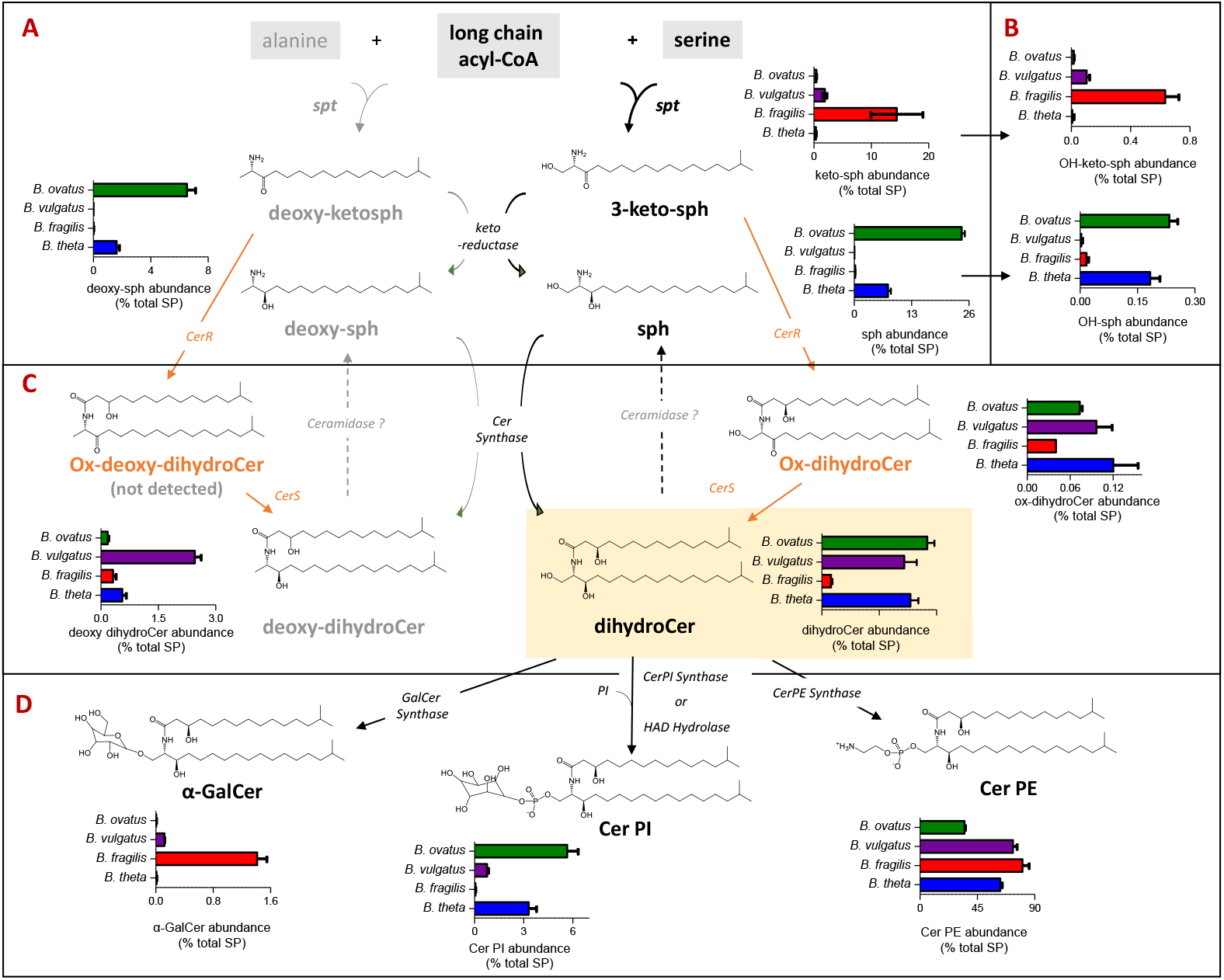
Dihydroceramide phosphoethanolamine (Cer PE) is the most abundant sphingolipid (SP) detected in all four *Bacteroides*. A) Relative abundance (expressed as % total PS) of sphingoid bases which include keto-sphinganines (keto-sph) and sphinganines (sph) derived from either serine or alanine (deoxy derivatives) among the four *Bacteroides* species (B) Relative abundance (expressed as % total SP) of hydroxy keto-sph and sph, exact position of hydroxylation is unknown. (C) Relative abundance (expressed as % total SP) of ceramide (Cer) lipids among the four *Bacteroides* species derived from either serine (dihydroCer) or alanine (deoxy-dihydroCer). (D) Relative abundance of complex *Bacteroides* SP (expressed as % total SP) including α-galactosyl dihydroCer (α-GalCer) and the phosphosphingolipids, dihydroCer phosphosethanolamines (Cer PE) and dihydroCer phosphoinositols (Cer PI). The biosynthesis of bacterial SP is initiated by serine palmitoyltransferase (SPT) which catalyses a reaction between a fatty acyl-CoA and serine, or alternatively alanine, to form keto-sph, or deoxy-keto-sph, respectively. The pathway to dihydroCer synthesis may proceed via that observed in eukaryotes i.e., via keto reductase activity (Lee *et al*., 2022) to form sph or alternatively bacterial Cer synthase (CerS) directly adds an acyl chain to 3-keto-sph producing an oxidised dihydroCer intermediate (ox-dihydroCer) which is then reduced to dihydroCer by bacterial Cer reductase (CerR) (Stankeviciute et al., 2021). DihydroCer represents the central hub of SP metabolism and undergoes modification with different head groups to produce the uniquely bacterial SP, Cer PE, Cer PI and α-GalCer (Brown *et al*., 2019; Wieland Brown *et al*., 2013). The biosynthesis of Cer PI involves either Cer PI synthase or haloalkanoate dehalogenase (HAD) hydrolase activity (Heaver. *et al*., 2022). *B. fragilis* α-GalCer was recently reported to be formed via a ceramide UDP-GalCer synthase (Okino et *al*., 2020)

Cer PE was identified as the most abundant SP in all 4 species with levels between 35% (*B. ovatus*) and 81% (*B. fragilis*) of the total SP detected (Figure 3D). In the current study Cer PI represented 3%, 6% and 1% of the total SP fraction in *B. thetaiotaomicron, B. ovatus*, and *B. vulgatus* respectively and this SP was not detected in *B. fragilis* (Figure 3D). Moreover α-GalCer, was detected in only *B. fragilis* and *B. vulgatus* (Figure 3D). Therefore, there is a wide range (both qualitatively and quantitatively) of SP and their intermediates produced across the *Bacteroides* species examined in this study.

### Plasmalogens and phosphoinositol (PI) lipids are not found in all *Bacteroides* species

DG, the simplest glycerol based membrane lipid and a metabolic intermediate to GP (Figure 2) accounts for between 3% (*B. fragilis*) and 13% (*B. ovatus*) of the total lipid content of *Bacteroides* (Figure 4A). Thereafter, the GP lipid fraction was dominated by diacyl PE (97% of total GP) in all four *Bacteroides* tested (Figure 4B) with minor amounts of lyso-PE (LPE) and diacyl PS also detected (Figure 4B). On the other hand diacyl PI was detected in lipid extracts of *B. thetaiotaomicron* and *B. ovatus* but not in *B. fragilis* and *B. vulgatus* (Figure 4B). Interestingly, we detected a number of plasmalogens (both diacylated (PE-P) and lyso-derivatives (LPE-P)) in *B. thetaiotaomicron* only (Figure 4B) at m/z [M-H]-of 576.4035, 590.4191, 604.4348, 618.4504, 632.4661, 646.4817, 660.4974 (PE-P) and 394.2364, 408.2521, 422.2677, 436.2834 (LPE-P) (Table S2). Fragments at m/z 196.0380 and 140.0118, corresponding to the loss of the plasmenyl group and the ethanolamine phosphate ion, respectively, were detected in negative mode MS2 spectra confirming that these plasmalogens are derived from PE. The presence of plasmalogens in *B. thetaiotaomicron* was also confirmed by acid hydrolysis (see Figure S2).

**Figure 4:**
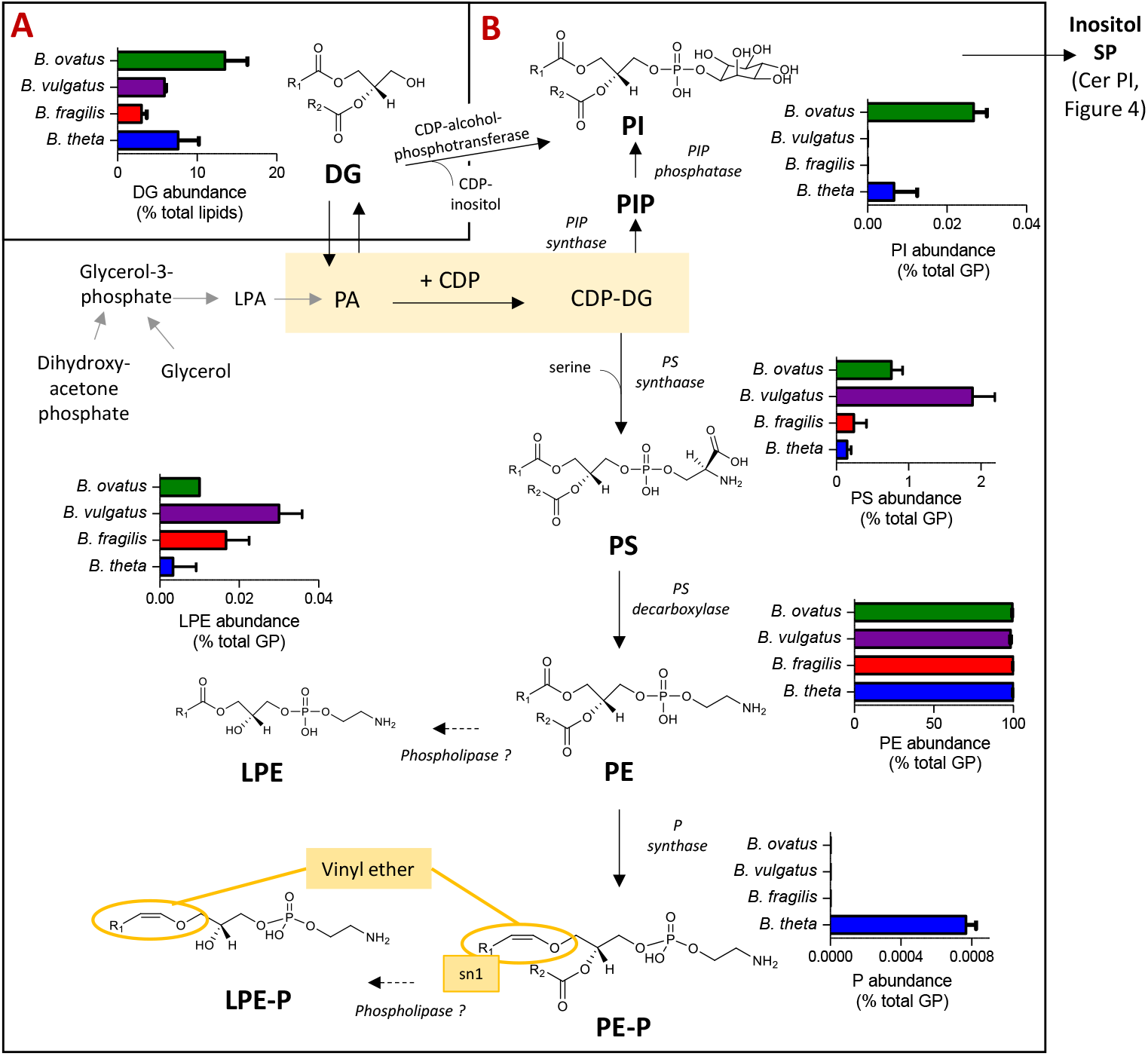
The presence of plasmalogens and phosphoinositol (PI) lipids is dependent on the *Bacteroides* species. A) Relative abundance (expressed as % total lipid) of diacylglycerols (DG) among the four *Bacteroides* tested. DG act as metabolic intermediates in the biosynthesis of *Bacteroides* glycerophospholipids (GP). (B) Relative abundance (expressed as % total GP) of glycerophosphoinositol (PI), glycerophosphoserine (PS), glycerophosphoethanolamines (PE), mono-acylated or lyso PE (LPE) and the vinyl-ether containing plasmalogens (P), PE-P and LPE-P. The pathways of *Bacteroides* derived GP shown are adapted from literature (Sohlenkamp & Geiger 2016; Zheng *et al*., 2019; Jackson *et al*., 2021; Heaver *et al*., 2022) Briefly, the biosynthesis of bacterial GP primarily start from the central metabolite cytidine diphosphate-DG (CDP-DG) leading to the production of PI in one direction or PS in the other. PI is formed via PI phosphate (PIP) intermediates (Sohlenkamp & Geiger 2016; Heaver *et al*., 2022) or directly via DG and CDP-alcohol-phosphotransferase (Heaver *et al*., 2022). PS acts as a metabolic intermediate and can undergo decarboxylation to form PE via PS decarboxylase. PE may also be formed from CDP-DG directly via a bifunctional Cardiolipin/PE synthase (not shown here) (Sohlenkamp & Geiger, 2016). In anaerobic bacteria, PE-P can be formed from PE via plasmalogen synthase (P synthase) (Jackson *et al*., 2021). In bacterial membranes, LPE are generated as metabolic intermediates in phospholipid synthesis or from membrane degradation via the action of phospholipases (Zheng *et al*., 2017). PA: glycerophosphatic acid; LPA: lyso PA

### A mutation in sphingolipid (SP) biosynthesis results in global changes in the lipid signature, including reductions in the levels of GlyL

Given that the synthesis of all lipids requires similar building blocks, we reasoned that disrupting the production of a class of lipids would result in major compensatory lipid signature alterations. To quantify these changes, we compared the lipid signatures of *B. thetaiotaomicron* wild-type with the Δ*SPT* mutant. The *SPT* gene encodes the enzyme required for the first step in SP biosynthesis, serine palmitoyltransferase. Multivariate analyses and comparison of lipid signatures showed that deletion of *SPT* results in dramatic changes to the lipid profile (Figure 6G). Moreover, these changes were not limited to SP lipids and we observed that 15 of the 26 lipid ‘subgroups’ or 86 of 170 individual lipids were significantly decreased in the *ΔSPT* mutant compared to the wild-type (>2 fold, *p* value (<0.05)). Not surprisingly, we observed that all of the SPs were depleted whilst GlyL and FL were also significantly decreased in the *ΔSPT* mutant relative to WT (Figure 6B). Moreover all PE (mono- and diacyl) and PE plasmalogens (mono- and diacyl) were also significantly reduced in the Δ*spt* mutant (Figure 6B). Indeed, there was close to a 60% overall decrease in the total identifiable lipids detected in the Δ*SPT* mutant relative to the WT (Figure S3B). Diacyl PI was the only lipid subgoup that increased in the Δ*SPT* mutant relative to WT (Figure 6C). Therefore a mutation in SP production results in global changes in the lipid profile of the membranes of *B. thetaiotaomicron*.

### A mutation in glycine lipid biosynthesis results in changes in the sphingolipid pool

The *glsB* gene encodes the enzyme responsible for the first step in glycine lipid biosythesis (Figure 2C) (Lynch *et al*, 2019). Therefore we compared the lipid signatures of *B. thetaiotaomicron* wild-type with the Δ*glsB* mutant. In total 8 of the 26 lipid ‘subgroups’ or 36 of the 170 individual lipids identified in this study were decreased in the Δ*glsB* mutant compared to WT (Figure 5A). As expected, GlyL and the related FL and complex ‘unknown’ lipids were depleted in the Δ*glsB* mutant extracts compared to WT (Figure 5B). On the other hand, 6 of the 26 lipid subgroups (or 37 of the 170 individual lipids) were significantly increased in the Δ*glsB* mutant compared to WT (Figure 5A). These lipids include SP subgroups Cer PE, dihydroCer and Cer PI (Figure 5C) and GP subgroups DG, diacyl PI and diacyl PS (Figure 5C). Therefore, it appears that depletion of the glycine lipids is compensated for by increasing other lipid groups, particularily SP and GP.

**Figure 5:**
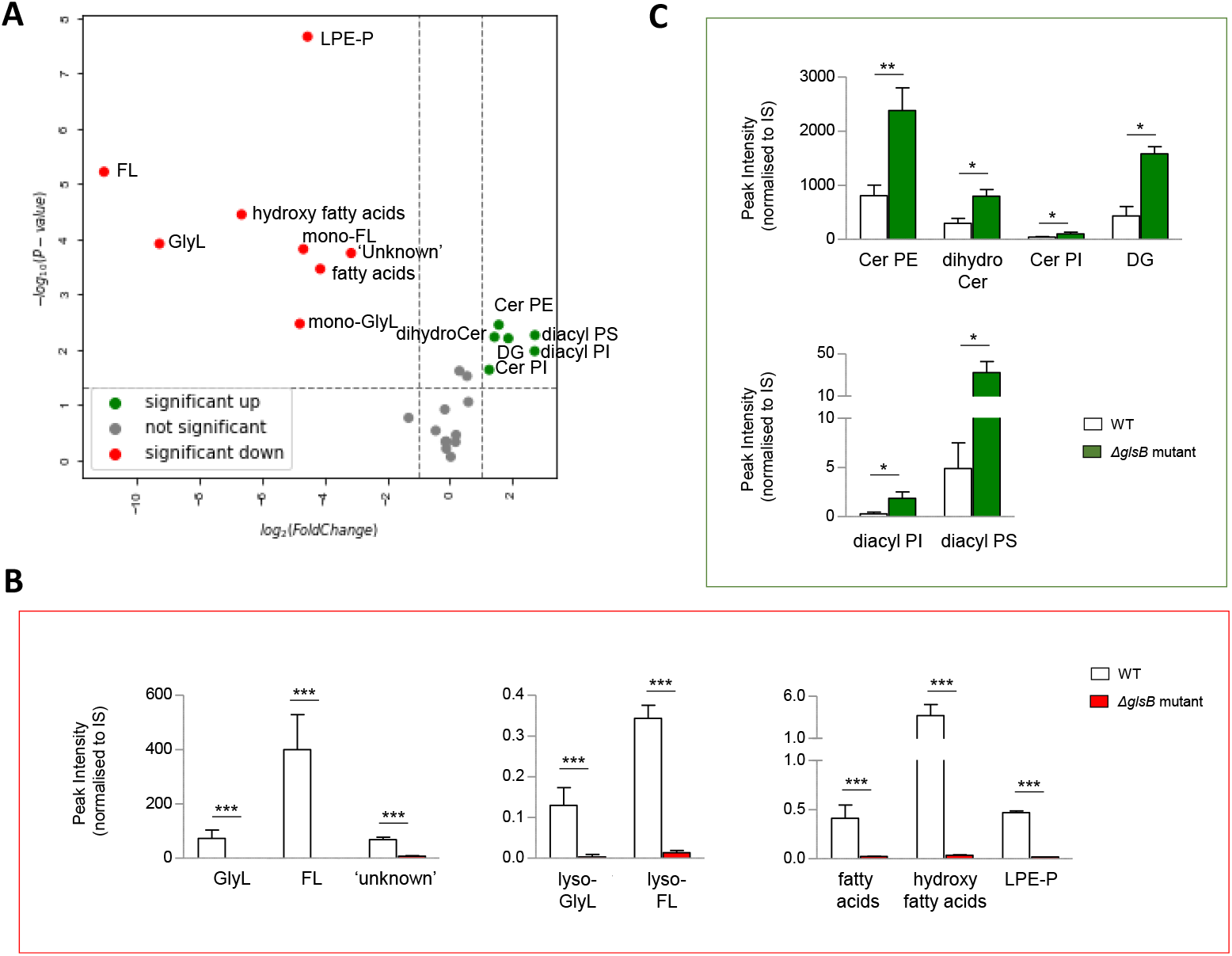
A mutation in glycine lipid biosynthesis (Δgls*B* mutant) results in changes in the sphingolipid (SP) pool relative to wild type (WT) *B. thetaiotaomicron*. (A) Univariate analyses (volcano plot) reveal significant (fold change >2 and t-test p-value < 0.05) changes to the lipid profile of *B. thetaiotaomicron* with 8 of the 26 lipid ‘subgroups’ significantly decreased in *B. thetaiotaomicron* Δgls*B* mutant versus WT, 6 significantly increased and 12 either unchanged or not detected (ND). (D) Glycine lipids (GlyL) and flavolipins (FL) were completely depleted in mutant extracts. Other lipids significantly decreased were a series of ‘unknown’ but predicted *N*-acyl amine lipids, fatty acids, hydroxy fatty acids and glycerophosphoethanolamines (PE) plasmalogens (P), (mono-acylated, LPE-P). (E) Many SP were significantly increased in mutant extracts versus wild-type and included dihydroceramides (dihydroCer), dihydroCer ethanolamines (Cer PE) dihydroCer inositols (Cer PI) Significant increases were also observed for diacylglycerols (DG), diacyl glycerophosphoinoistols (PI) and diacyl glycerophosphoserines (PS). Note: All data shown are mean values of n=3 ± SD. *** P < 0.001, ** P<0.005, * P<0.05 and was log transformed prior to univariate analyses (data shown is un-transformed).

## Discussion

Microbial lipids are becoming recognised as interkingdom signalling molecules and, as such, they represent an interesting avenue for the potential development of new and novel biomarkers and/or therapeutics. In this study we set out to construct a lipid map for several mammalian gut resident *Bacteroides*, namely *B. thetaiotaomicron, B. fragillis, B*.*ovatus* and *B. vulgatus*. Individually, these selected *Bacteroides* may account for up to 6% of intestinal bacteria in healthy humans with *B. vulgatus* reported to represent up to 40% in patients with Crohn’s Disease (Wang *et al*., 2021).

In this study we have identified 170 individual lipids, representing 4 LIPID MAPS categories and 26 lipid ‘subgroups’ (Table S2), across the 4 selected species. PE (diacylated) was noted as the major core lipid subgroup in all four *Bacteroides* (Figure 4). Our data suggests that PE is likely formed via the decarboxylation of PS given that PS (diacylated) was also detected, albeit in relatively minor amounts, in all four species. Different lyso-PE (LPE) species were also noted as minor core lipids in all four *Bacteroides* (Figure 4). These lipids are likely generated as metabolic intermediates in PE synthesis or from the degradation of PE (Zheng *et al*. 2017). The role of LPE in bacteria remains poorly characterised although it is believed that LPE may mitigate membrane stress induced by non-bilayer lipids such as cardiolipin. Whilst CL is an important GP in some Gram negative bacteria such as *E. coli* (Sohlenkamp & Geiger, 2016) and CL has been reported in some *Bacteroides* (Rizza *et al*.,1970, Wardle *et al*.,1996) we did not detect CL in any of the *Bacteroides* used in this study.

This study is the first to detect PE plasmalogens, both mono- and diacyl, in *Bacteroides*. Interestingly we only detected plasmalogens in *B. thetaiotaomicron* (Figure 4) suggesting that these lipids are not ubiquitous in this genus. Plasmalogens are widely distributed in eukaryotes where they are important for membrane structure, membrane trafficking and cell signalling (Dean & Lodhi, 2018). Reduced levels of circulating plasmalogens has been linked to several metabolic and neurological diseases including diabetes and Alzheimers disease (Dean & Lodhi, 2018; Su *et al*., 2019).

We previously described a family of *N*-acyl amines in *B. thetaiotaomicron* called glycine lipids (Lynch *et al*., 2019). In this study we have confirmed the presence of glycine lipids in all species of *Bacteroides* tested suggesting that these lipids are widespread in this genus (Figure 2). Indeed bioinformatic analysis indicates that the genetic potential for the production of glycine lipids is restricted to genera in the phylum Bacteroidota (our unpublished data). Nonetheless there were some differences in the quantities of specific glycine lipids although the physiological relevance of these differences is not clear. FL-654 was the most abundant glycine lipid detected in all *Bacteroides* tested. FL-654 has, in several studies, been reported to act as a TLR2 agonist suggesting a possible role in inflammation in the host (Clark *et al*., 2013; Nemati *et al*., 2017; Nichols *et al*., 2020a). A recent report shows that the chronic (7-week) intraperitoneal administration of FL-654 to high fat diet (HFD)-fed low-density lipiproprotein receptor (Ldlr-/-) mice lowered cholesterol, attenuated atherosclerosis progression, and decreased markers of liver injury compared with vehicle control-injected mice (Millar *et al*., 2022). The glycine lipid family also includes high-molecular lipid molecules such as Lipid 1256 that consist of a diacyl glycerophosphoglycerol (PG) linked to the FL (Nichols *et al*., 2020b). Lipid 1256 was reported to be an even more potent TLR2 ligand than the related GlyL and FL and has been studied in *Porphyromonas*, a relative of *Bacteroides* generally found in the oral cavity (Nichols *et al*., 2020b). In the current study, we initially assumed that Unknown_1259 (Table S2) was the same as Lipid 1256 but given the absence of PG in our *Bacteroides* extracts and the difference of a proton in the observed m/z we believe that Unknown_1259 is more likely to be FL linked to PE. Work is currently been carried out to confirm the structure of this potentially novel lipid.

Compared to well-studied bacteria such as *E. coli*, the membranes of *Bacteroides* do have an unusual lipid composition in that approximately, 50% of the lipids that are extractable with chloroform-methanol are SP or free ceramides (Salyers, 1984). In the present study, SP represented between 19% (*B. ovatus*) and 29% (*B. vulgatus*) of the total lipids detected by isopropanol extraction. For the most part, dihydroCer and Cer PE represent the core SP (Figure 3) detected and this is typical of bacteria in the phylum *Bacteroidota* (Panevska *et al*., 2019). Both dihydroCer and Cer PE have been shown to be negatively correlated with inflammation and Inflammatory Bowel Disese (IBD) in humans (Brown *et al*., 2019). The same authors also reported the detection of deoxy-dihydroCer in *B. thetaiotaomicron* formed via the utilisation of alanine, rather than serine, by SPT (Figure 6). In the present study, putative dexoy-dihydroCer was detected in the four *Bacteorides* tested, most notably higher in *B. vulgatus* (Figure 3). In addition, a putative oxidised dihydroCer (ox-dihydroCer) lipid was detected as a minor core lipid supporting the notion that dihydroCer synthesis may proceed via bacterial ceramide synthase (CerS) which directly adds an acyl chain to keto-sph producing ox-dihydroCer which is then reduced to dihydroCer by bacterial ceramide reductase (CerR) (Stankeviciute *et al*., 2021, Figure 3). The upstream SP, keto-sph was particularily abundant in *B. fragilis* suggesting a potentially slower conversion to dihydroCer and/or faster rate of sythesis via SPT (see Figure 3A). This highlights potentially important species-specific differences in enzyme kinetics and flux through the SP biosynthetic pathway. The presence of sph and deoxy-sph in *B. thetaiotaomicron* and *B. ovatus* may be indicative of increased keto-reductase activity (Lee *et al*., 2022) and/or a slower rate of conversion to dihydroCer in these *Bacteroides* species. It may also be interpreted to indicate the presence of a ceramidase, absent in *B. fragilis* and *B. vulgatus*, that hydrolyses dihydroCer to sph (Stankevicute *et al*., 2021).

**Figure 6:**
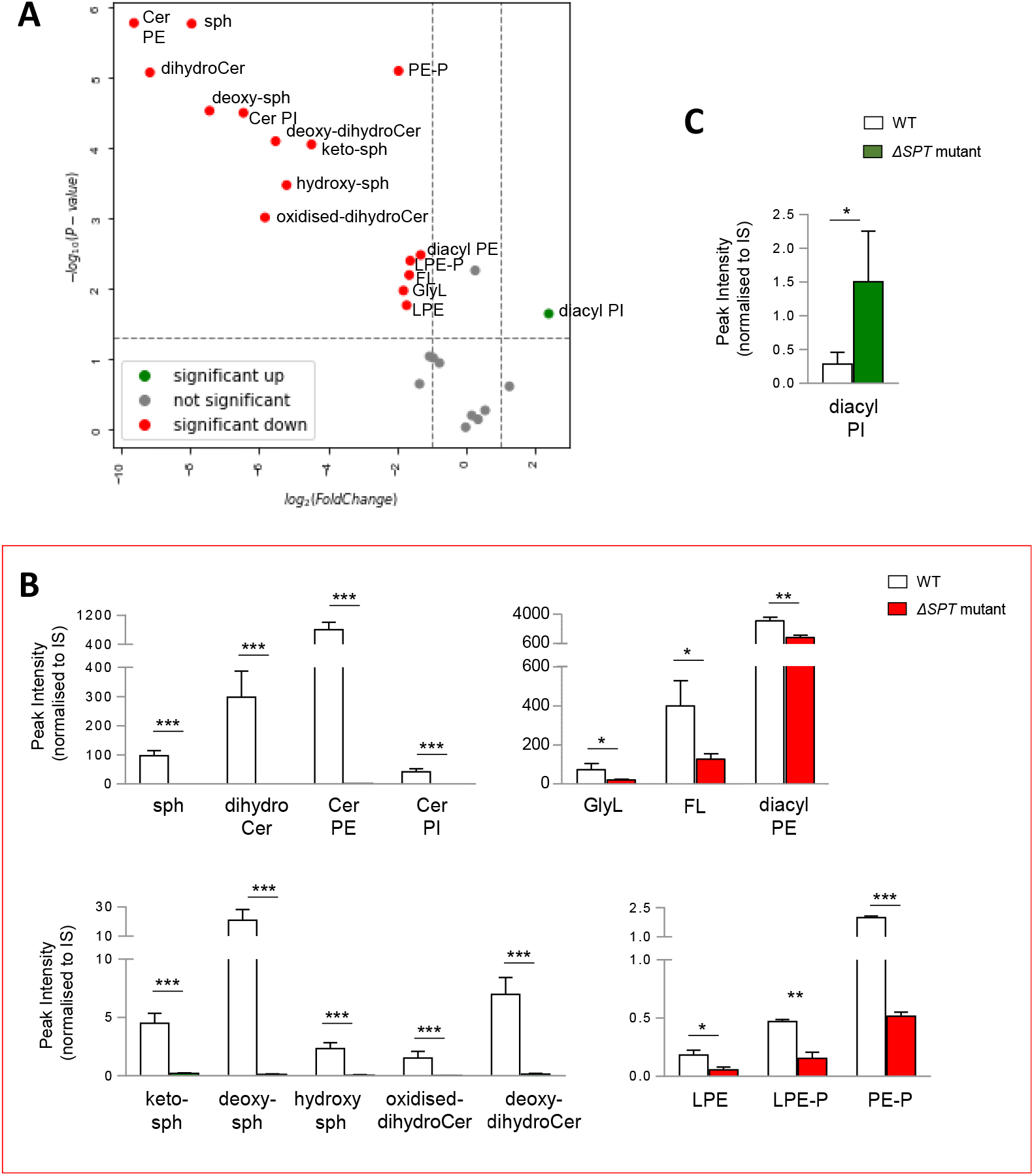
A mutation in sphingolipid (SP) biosynthesis (ΔSPT mutant) results in significant changes in the lipid signature of *B. thetaiotaomicron* relative to wild type (WT) (A) Univariate analyses (volcano plot) reveal significant (fold change >2 and t-test p-value < 0.05) changes to the lipid profile of *B. thetaiotaomicron* whereby 15 of the 26 lipid ‘subgroups’ were significantly down (red) in *B. thetaiotaomicron* Δ*SPT* mutant versus WT, 1 significantly up (green) and 10 either unchanged or not detected (grey) (B)The SP ‘subgroups’ detected in *B. thetaiotaomicron* were completely depleted in mutant extracts. Other lipids significantly decreased were glycine lipids (GlyL) and flavolipins (FL) as well as both lyso-(L) and di- (diacyl) acylated glycerophosphoethanolamines (PE), and their respective plasmalogens (P) (C) Diacyl glycerophosphoinoistols (PI) were the only lipid subgroup to be significantly decreased in mutant versus wt. Note: All data shown are mean values of n=3 ± SD. *** P < 0.001, ** P<0.005, * P<0.05 and was log transformed prior to univariate analyses (data shown is un-transformed).

Depending on the species, two other complex SP were detected in the present study. Cer PI, which consist of a inositol phosphate on a sphingoid backbone, were detected in *B. thetaiotaomicron* and *B*.*ovatus* as reported previously by Brown *et al*. (2019). However, we also show that Cer PI are present in *B. vulgatus* (Figure 3). The gene clusters reponsible for inositol lipid synthesis in *Bacteroides* have recently been described (Brown *et al*., 2019; Heaver *et al*., 2022; Sartorio *et al*, 2022;) and involves either PI Cer synthase, typical of *B. thetaiotaomicron* and *B*.*ovatus* or HAD hydrolase activity, typical of *B. vulgatus* (Figure 3). Heaver *et al*. (2022) report that inositol and inositol lipids are likely implicated in resistance to host immune defences through their roles in the structure of the membrane and the capsule, and important for fitness in the mammalian gut. The ‘non-phosphate’ containing complex glycosphingolipids, α-GalCer, were detected in *B. fragilis* and, to a lesser extent in *B. vulgatus*. This is consistent with previous reports on the generation of α-GalCer by *B. fragilis* (Wieland-Brown *et al*., 2013; von Greichten *et al*., 2017) and lower levels by *B. vulgatus* (von Greichten *et al*., 2017). *B. fragilis* α-GalCer recently reported to be formed via a ceramide UDP-galactosylceramide synthase (Okino *et al*., 2020) has been shown to be potent stimulators for invariant NKT cells (Wieland-Brown *et al*., 2013) whereby the sphinganine chain branching is a critical determinant of NKT activation (Oh *et al*., 2021).

SP and glycine lipids are found in all Bacteroides tested and are likley to have a structural role in the membrane of these bacteria. Therefore we examined the changes in lipid signatures following mutations to these two major bioactive lipiid pathways in *B. thetaiotaomicron*. Using a mutation in the glsB gene we showed that *B. thetaiotaomicron* compensate for the absence of glycine lipids by increasing some SP, DG, diacyl PI and diacyl PS. In contrast there was an overall decrease in lipid diversity in the ΔSPT mutant, including a significant decrease in many glycine lipids, PE (both diacyl and lyso) and PE plasmalogens (both diacyl and lyso). The only attempt at compensation appears to be the signifcant increase in PI (Figure 6). Given that Cer PI are not formed in the ΔSPT mutant, the increase in PI may simply be due to their reduced coupling to ceramide (Figure 3D). Thus, perturbations in glycine lipid or SP biosynthesis results in significant, yet distinct, changes in the levels of other lipids suggesting the existence of compensatory mechanisms required to maintain the functionality of the bacterial membrane. The relatively dramatic global lipid decreases in the ΔSPT mutant may suggest that SP have a key structural role in the membranes of *Bacteroides* i.e. without SP other key membrane lipids cannot organise/insert in the membrane and are therefore depleted from lipidome.

In summary, *Bacteroides* produce diverse lipids, some of which are species dependent e.g. plasmalogen production in *B. thetaiotaomicron* (Figure 4). The exact role of these plasmalogens, in or between bacteria, or as molecular signalling molecules to the host remains to be elucidated but the generation of plasmalogens by important gut bacteria provides a potential opportunity to modulate plasmalogen production as a therapeutic strategy (Paul *et al*., 2019). For *B. fragilis*, the most noticable difference in the lipid profile was the relatively higher abundance of lyso-FL (Figure 2), keto-sph, and α-GalCer (Figure 3). The bioactivity of the latter has received considerable attention to date (Wieland-Brown *et al*., 2013; Oh *et al*., 2021) and certain lyso-FL species have been shown to have TLR2 agonist activity (Clark *et al*., 2013). However little is known on the bio-activity of bacterially derived keto-sph. The most notable difference in the lipid signatures of *B. ovatus*, relative to the other species was the accumulation of more DG, PI (Figure 4) and Cer PI (Figure 3). *B. ovatus* ATCC8483 has been shown to reduce mucosal inflammation by up-regulating IL-22 secretion (Ihekweazu *et al*., 2019, 2021). *B. vulgatus* had relatively more ‘unknown’ lipids (Figure 2), LPE and PS (Figure 4), and deoxy-dihydroCer. Given the association of *B*.*vulgatus* with IBD (Wang *et al*., 2021), these observations may prove important. The ‘unknown’ lipids are structurally related to Lipid 1256, a potent TLR2 ligand which promotes the production of pro-inflammatory cytokines. In addition, deoxy-dihydroCer are ‘dead-end’ toixc lipids (Duan & Merrill, 2015) that are implicated in the progression of type 2 Diabetes (Sen *et al*., 2021; Hannich *et al*., 2021). In short, this lipid profilling data will provide a useful platform for the further characterisation of the lipid-based host-microbe dialogue and the influence of microbial lipids on host health and disease states.

## Acknowlegements

We sincerely thank Eric Brown for providing the ΔSPT mutant and Eric Martens for Bacteroides isolates. This work was conceived and designed by ER, DJC and SAJ. The experimental work was performed by ER (MS) and BGP (bacteria). ER, DJC and SAJ analysed the data. ER, LAG, DJC and SAJ wrote and edited the manuscript. This work was funded by EU H2020-MSCA-No. 887019 “OMIT” and APC CoFund Apex award APEX No.754535 to ER, SAJ. BGP, DJC, SAJ are supported by Science Foundation Ireland Centres Grant SFI/12/RC/2273 P2 to APC Microbiome Ireland. SAJ is also funded by SFI: EU Joint Programme Initiative CABALA for Health No. 3358 and Ireland Department of Agriculture, Food and the Marine (DAFM) Award No. DAFM 17-RD-US-ROI.

## Supplementary Information

### Supporting Information

**Table SI.** Chromatographic gradient used for lipid profiling

| Time (min) | % A | % B | Curve |
| --- | --- | --- | --- |
| Initial | 60 | 40 | Initial |
| 2.0 | 57 | 43 | 6 |
| 2.1 | 50 | 50 | 1 |
| 12.0 | 46 | 54 | 6 |
| 12.1 | 30 | 70 | 1 |
| 18.0 | 1 | 99 | 6 |
| 18.1 | 60 | 40 | 6 |
| 20 | 60 | 40 | 1 |

**Figure S1:**
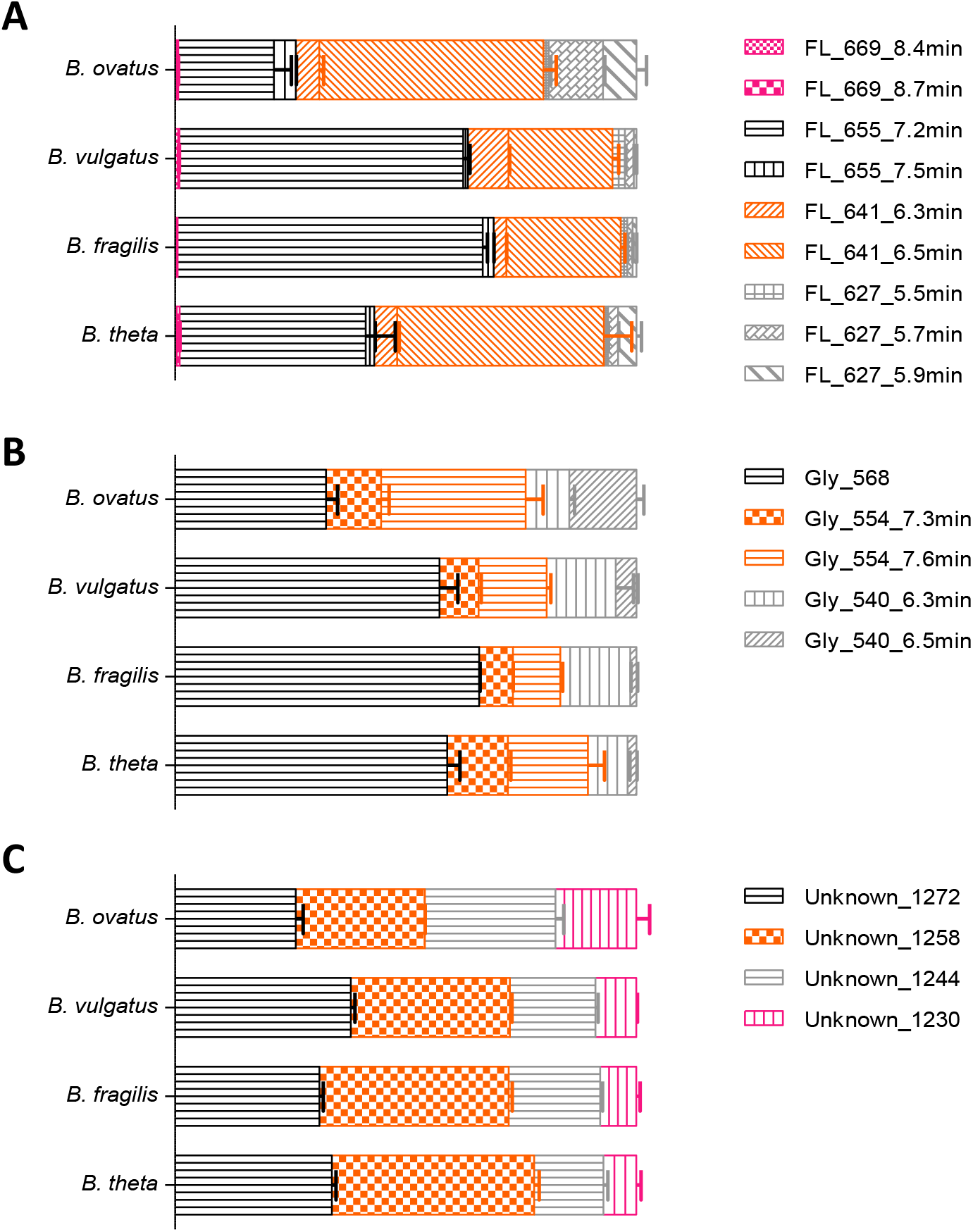
Relative abundance of each Flavolipin (FL, A), glycine lipid (GLyL, B) and ‘unknown’ lipid (C) in each Bacteroides species

**Figure S2.**
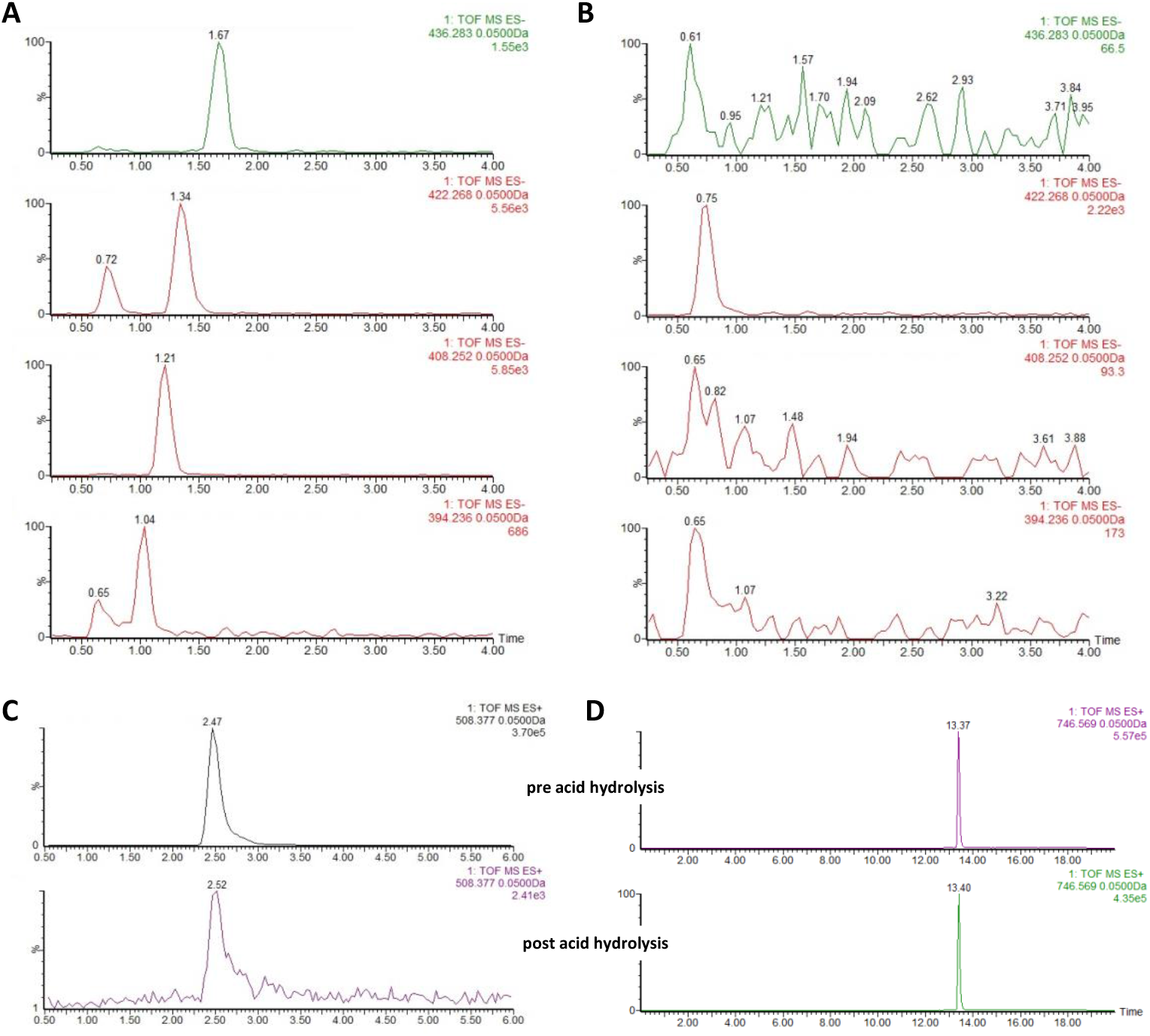
The presence of plasmalogens is confirmed in *B. thetaiotaomicron* extracts. (A) Presence of lyso-PE (LPE) plasmalogens (P) in *B. thetaiotaomicron* wild-type extracts at t_R_ 1.04 (LPE (P-13:0)), 1.21 (LPE (P-14:0)), 1.34 (LPE (P-15:0)) and 1.67 (LPE (P-16:0)) mins in the absence of acid hydrolysis (B) Absence of LPE-P in B. thetaiotaomicron wild-type extracts subjected to acid hydrolysis. (C) A positive control, lyso-glycerophosphocholine (LPC) P-18:0 was employed in acid hydrolysis experiments, in the absence of acid it was detected at 2.47 min and when subjected to acid hydrolysis LPE (P-18:0) was greatly reduced. A negative control, diacyl PE (PE 18:0/18:1) was employed in acid hydrolysis experiments, in the absence of acid it was detected at 2.4 min and when subjected to acid hydrolysis PE (18:0/18:1) was largely unaffected.

**Figure S3.**
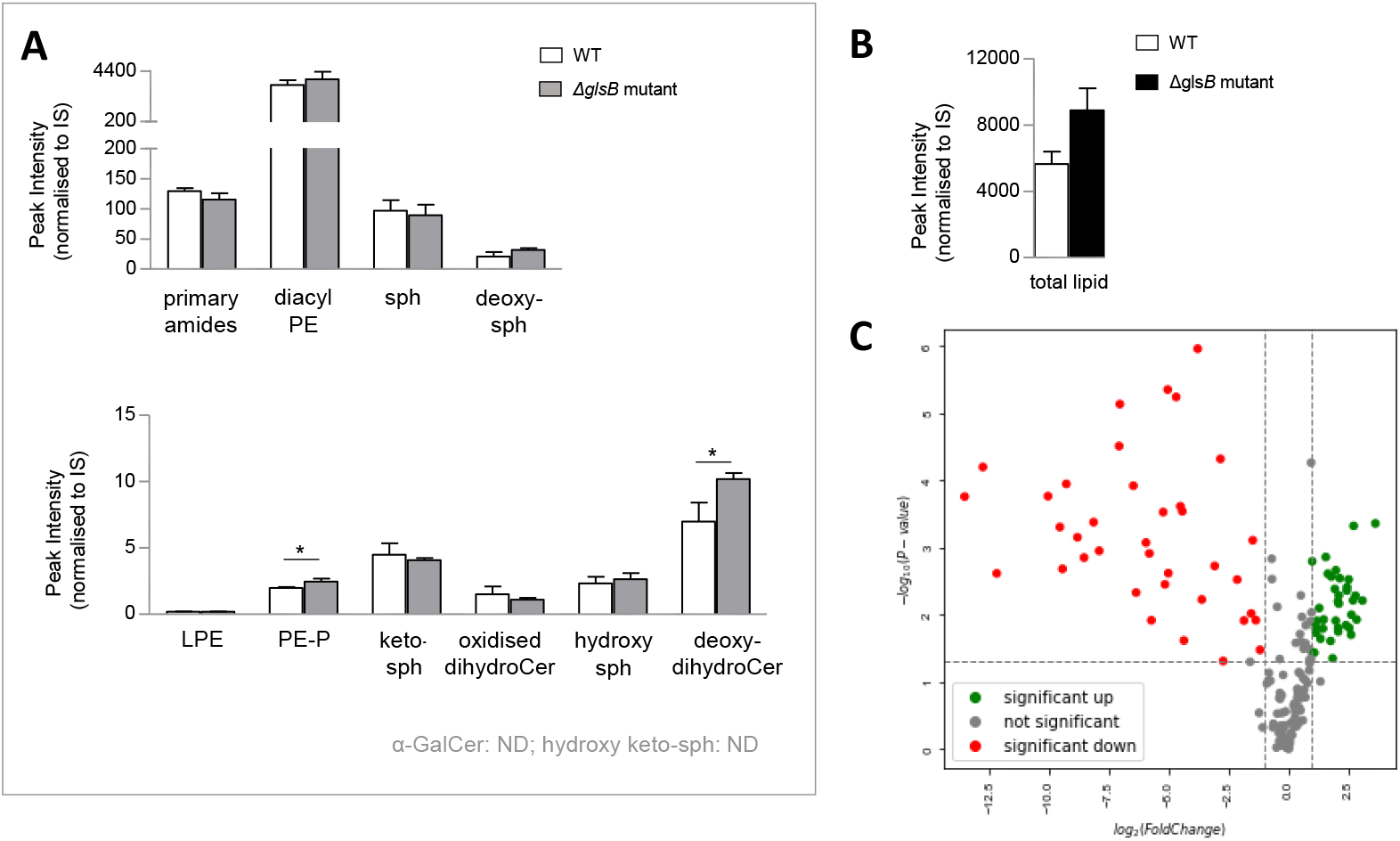
Effect of *glsB* mutation the lipid profile of *B. thetaiotaomicron* (A) 12 of the 26 lipid ‘subgroups’ were either not significantly altered or not detected (ND) in *B. thetaiotaomicron Δ*gls*B* mutant versus wild-type (WT) (B) Mean total lipid profile (normalised to internal standard) of identified lipids in Δ*glsB* mutant versus WT extracts (C) Univariate analyses (volcano plot) reveal significant (fold change >2 and t-test p-value < 0.05) changes whereby 36 lipids were significantly down (red) in *B. thetaiotaomicron ΔglsB* mutant versus wild-type,37 were significantly up (green) and 97 were either unchanged or not detected (grey). Note: All data shown are mean values of n=3 ± SD. *** P < 0.001, ** P<0.005, * P<0.05 and was log transformed prior o univariate analyses (data shown is un-transformed).

**Figure S4.**
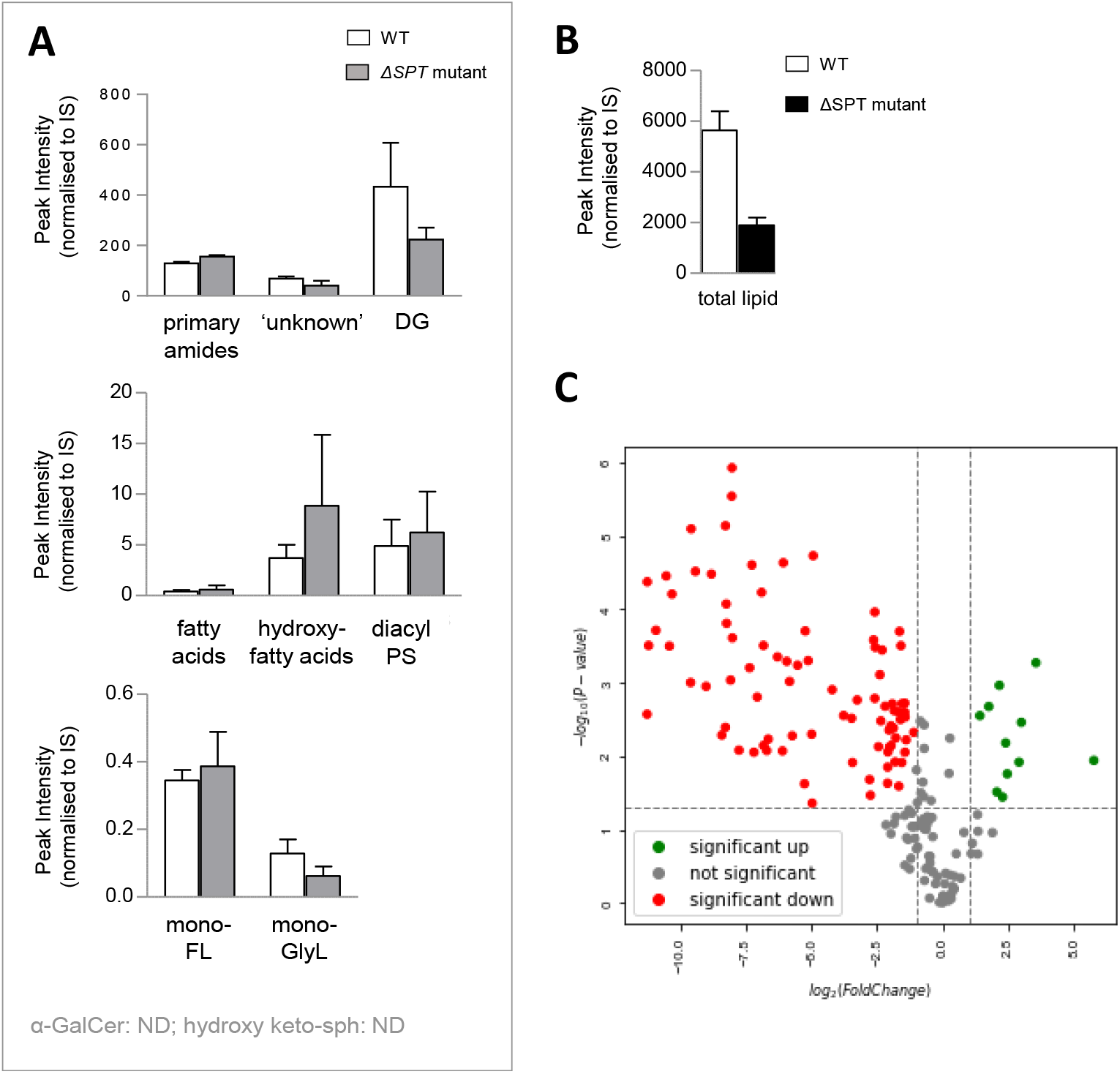
Effect of *SPT* mutation on lipid profile of *B. thetaiotaomicron* (A) 10 of the 26 lipid subgroups were not significantly altered (as determined by both fold change > 2 and p-value <0.05) or not detected in *B. thetaiotaomicron ΔSPT* mutant versus wild-type (WT). (B) Mean total lipid profile (normalised to internal standard) of identified lipids in *ΔSPT* mutant versus WT extracts (C) Univariate analyses (volcano plot) reveal significant (fold change >2 and t-test p-value < 0.05) changes to the lipid profile of *B. thetaiotaomicron* whereby 86 of the 170 lipids were significantly were significantly down (red) in *B. thetaiotaomicron ΔSPT* mutant versus WT, 11 were significantly up (green) and 73 were either unchanged or not detected (grey). All data shown are mean values of n=3 ± SD. *** P < 0.001, ** P<0.005, P<0.05 and was log transformed prior to univariate analyses (data shown is un-transformed).

**Table S2.** Lipids detected by LC-MS in Bacteroide isopropanol extracts

| Lipid | Formula | Ion | m/z | t<sub>R</sub> | Lipid Category | Lipid Class | Lipid Subclass |
| --- | --- | --- | --- | --- | --- | --- | --- |
| DG (25:0) | C28H54O5 | M+Na | 493.3869 | 5.00 | Glycerolipids (GL) | Diradylglycerols | Diacylglycerols (DG) |
| DG (26:0) | C29H56O5 | M+Na | 507.4025 | 5.50 | Glycerolipids (GL) | Diradylglycerols | Diacylglycerols (DG) |
| DG (27:0) | C30H58O5 | M+Na | 521.4182 | 6.60 | Glycerolipids (GL) | Diradylglycerols | Diacylglycerols (DG) |
| DG (28:0)_7.3min | C31H60O5 | M+Na | 535.4338 | 7.26 | Glycerolipids (GL) | Diradylglycerols | Diacylglycerols (DG) |
| DG (28:0)_7.6min | C31H60O5 | M+Na | 535.4338 | 7.62 | Glycerolipids (GL) | Diradylglycerols | Diacylglycerols (DG) |
| DG (28:0)_8.0min | C31H60O5 | M+Na | 535.4338 | 7.94 | Glycerolipids (GL) | Diradylglycerols | Diacylglycerols (DG) |
| DG (29:0)_8.4min | C32H62O5 | M+Na | 549.4495 | 8.39 | Glycerolipids (GL) | Diradylglycerols | Diacylglycerols (DG) |
| DG (29:0)_8.7min | C32H62O5 | M+Na | 549.4495 | 8.74 | Glycerolipids (GL) | Diradylglycerols | Diacylglycerols (DG) |
| DG (30:0)_9.6min | C33H64O5 | M+Na | 563.4651 | 9.63 | Glycerolipids (GL) | Diradylglycerols | Diacylglycerols (DG) |
| DG (30:0)_10.1min | C33H64O5 | M+Na | 563.4651 | 10.12 | Glycerolipids (GL) | Diradylglycerols | Diacylglycerols (DG) |
| DG (30:0)_10.7min | C33H64O5 | M+Na | 563.4651 | 10.64 | Glycerolipids (GL) | Diradylglycerols | Diacylglycerols (DG) |
| DG (31:0)_11.2min | C34H66O5 | M+Na | 577.4808 | 11.24 | Glycerolipids (GL) | Diradylglycerols | Diacylglycerols (DG) |
| DG (31:0)_11.6min | C34H66O5 | M+Na | 577.4808 | 11.63 | Glycerolipids (GL) | Diradylglycerols | Diacylglycerols (DG) |
| DG (32:0)_12.8min | C35H68O5 | M+Na | 591.4968 | 12.80 | Glycerolipids (GL) | Diradylglycerols | Diacylglycerols (DG) |
| DG (32:0)_13.0min | C35H68O5 | M+Na | 591.4968 | 13.04 | Glycerolipids (GL) | Diradylglycerols | Diacylglycerols (DG) |
| DG (32:1) | C35H66O5 | M+Na | 589.4811 | 10.94 | Glycerolipids (GL) | Diradylglycerols | Diacylglycerols (DG) |
| DG (33:0) | C36H70O5 | M+Na | 605.5121 | 13.28 | Glycerolipids (GL) | Diradylglycerols | Diacylglycerols (DG) |
| DG (34:0) | C37H72O5 | M+Na | 619.5277 | 13.67 | Glycerolipids (GL) | Diradylglycerols | Diacylglycerols (DG) |
| DG (34:1) | C37H70O5 | M+Na | 617.5124 | 13.10 | Glycerolipids (GL) | Diradylglycerols | Diacylglycerols (DG) |
| C13:0 | C13H26O2 | M-H- | 213.1860 | 1.57 | Fatty Acyls (FA) | Fatty Acids & Conjugates | Straight/Branched |
| C14:0 | C14H28O2 | M-H- | 227.2018 | 1.94 | Fatty Acyls (FA) | Fatty Acids & Conjugates | Straight/Branched |
| C15:0 | C15H30O2 | M-H- | 241.2175 | 2.20 | Fatty Acyls (FA) | Fatty Acids & Conjugates | Straight/Branched |

**Table S2.** Continued
| Lipid | Formula | Ion | m/z | t<sub>R</sub> | Lipid Category | Lipid Class | Lipid Subclass |
| --- | --- | --- | --- | --- | --- | --- | --- |
| C16:0 | C16H32O2 | M-H- | 255.2331 | 2.79 | Fatty Acyls (FA) | Fatty Acids & Conjugates | Straight/Branched |
| hydroxy C15:0 | C15H30O3 | M-H- | 257.2122 | 1.38 | Fatty Acyls (FA) | Fatty Acids & Conjugates | Hydroxy Fatty Acids |
| hydroxy C16:0 | C16H32O3 | M-H- | 271.2279 | 1.63 | Fatty Acyls (FA) | Fatty Acids & Conjugates | Hydroxy Fatty Acids |
| hydroxy C17:0 | C17H34O3 | M-H- | 285.2435 | 1.84 | Fatty Acyls (FA) | Fatty Acids & Conjugates | Hydroxy Fatty Acids |
| C16:0 amide | C16H33NO | M+H+ | 256.2635 | 1.87 | Fatty Acyls (FA) | Fatty Amides | Primary amides |
| C18:0 amide | C18H37NO | M+H+ | 284.2948 | 2.72 | Fatty Acyls (FA) | Fatty Amides | Primary amides |
| FL_669_8.4min | C38H72N2O7 | M+H+ | 669.5412 | 8.38 | Fatty Acyls (FA) | Fatty Amides | N-acyl amines |
| FL_669_8.7min | C38H72N2O7 | M+H+ | 669.5412 | 8.67 | Fatty Acyls (FA) | Fatty Amides | N-acyl amines |
| FL_655_7.2min | C37H70N2O7 | M+H+ | 655.5256 | 7.22 | Fatty Acyls (FA) | Fatty Amides | N-acyl amines |
| FL_655_7.5min | C37H70N2O7 | M+H+ | 655.5256 | 7.50 | Fatty Acyls (FA) | Fatty Amides | N-acyl amines |
| FL_641_6.3min | C36H68N2O7 | M+H+ | 641.5099 | 6.28 | Fatty Acyls (FA) | Fatty Amides | N-acyl amines |
| FL_641_6.5min | C36H68N2O7 | M+H+ | 641.5099 | 6.53 | Fatty Acyls (FA) | Fatty Amides | N-acyl amines |
| FL_627_5.5min | C35H66N2O7 | M+H+ | 627.4943 | 5.48 | Fatty Acyls (FA) | Fatty Amides | N-acyl amines |
| FL_627_5.7min | C35H66N2O7 | M+H+ | 627.4943 | 5.68 | Fatty Acyls (FA) | Fatty Amides | N-acyl amines |
| FL_627_5.9min | C35H66N2O7 | M+H+ | 627.4943 | 5.92 | Fatty Acyls (FA) | Fatty Amides | N-acyl amines |
| mono_FL_431 | C22H42N2O6 | M+H+ | 431.3116 | 1.28 | Fatty Acyls (FA) | Fatty Amides | N-acyl amines |
| mono_FL_417 | C21H40N2O6 | M+H+ | 417.2959 | 1.14 | Fatty Acyls (FA) | Fatty Amides | N-acyl amines |
| GlyL_568 | C34H65NO5 | M+H+ | 568.4935 | 8.44 | Fatty Acyls (FA) | Fatty Amides | N-acyl amines |
| GlyL_554_7.3min | C33H63NO5 | M+H+ | 554.4784 | 7.26 | Fatty Acyls (FA) | Fatty Amides | N-acyl amines |
| GlyL_554_7.6min | C33H63NO5 | M+H+ | 554.4784 | 7.56 | Fatty Acyls (FA) | Fatty Amides | N-acyl amines |
| GlyL_540_6.3min | C32H61NO5 | M+H+ | 540.4628 | 6.34 | Fatty Acyls (FA) | Fatty Amides | N-acyl amines |
| GlyL_540_6.5min | C32H61NO5 | M+H+ | 540.4628 | 6.53 | Fatty Acyls (FA) | Fatty Amides | N-acyl amines |
| mono_ GlyL_330 | C18H35NO4 | M+H+ | 330.2639 | 1.31 | Fatty Acyls (FA) | Fatty Amides | N-acyl amines |
| Unknown_1273 | C70H134N3O14P | M+H+ | 1272.9676 | 14.51 | Fatty Acyls (FA) | Fatty Amides | N-acyl amines |
| Unknown_1259 | C69H132N3O14P | M+H+ | 1258.9520 | 14.49 | Fatty Acyls (FA) | Fatty Amides | N-acyl amines |
| Unknown_1245 | C68H130N3O14P | M+H+ | 1244.9363 | 14.42 | Fatty Acyls (FA) | Fatty Amides | N-acyl amines |

**Table S2.** Continued
| Lipid | Formula | Ion | m/z | t<sub>R</sub> | Lipid Category | Lipid Class | Lipid Subclass |
| --- | --- | --- | --- | --- | --- | --- | --- |
| Unknown_1231 | C67H128N3O14P | M+H+ | 1230.9207 | 14.39 | Fatty Acyls (FA) | Fatty Amides | N-acyl amines |
| PE (25:0) | C30H60NO8P | M+H+ | 594.4129 | 3.42 | Glycerophospholipids (GP) | Glycerophosphoethanolamines (PE) | Diacyl PE |
| PE (26:0)_3.6min | C31H62NO8P | M+H+ | 608.4286 | 3.64 | Glycerophospholipids (GP) | Glycerophosphoethanolamines (PE) | Diacyl PE |
| PE (26:0)_3.8min | C31H62NO8P | M+H+ | 608.4286 | 3.78 | Glycerophospholipids (GP) | Glycerophosphoethanolamines (PE) | Diacyl PE |
| PE (26:0)_3.9min | C31H62NO8P | M+H+ | 608.4286 | 3.91 | Glycerophospholipids (GP) | Glycerophosphoethanolamines (PE) | Diacyl PE |
| PE (26:1) * | C31H60NO8P | M+H+ | 606.4129 | 3.3 | Glycerophospholipids (GP) | Glycerophosphoethanolamines (PE) | Diacyl PE |
| PE (27:0)_4.2min | C32H64NO8P | M+H+ | 622.4442 | 4.18 | Glycerophospholipids (GP) | Glycerophosphoethanolamines (PE) | Diacyl PE |
| PE (27:0)_4.3min | C32H64NO8P | M+H+ | 622.4442 | 4.27 | Glycerophospholipids (GP) | Glycerophosphoethanolamines (PE) | Diacyl PE |
| PE (27:1) * | C32H62NO8P | M+H+ | 620.4286 | 3.5 | Glycerophospholipids (GP) | Glycerophosphoethanolamines (PE) | Diacyl PE |
| PE (28:0)_4.7min | C33H66NO8P | M+H+ | 636.4599 | 4.68 | Glycerophospholipids (GP) | Glycerophosphoethanolamines (PE) | Diacyl PE |
| PE (28:0)_4.9min | C33H66NO8P | M+H+ | 636.4599 | 4.89 | Glycerophospholipids (GP) | Glycerophosphoethanolamines (PE) | Diacyl PE |
| PE (28:0)_5.1min | C33H66NO8P | M+H+ | 636.4599 | 5.10 | Glycerophospholipids (GP) | Glycerophosphoethanolamines (PE) | Diacyl PE |
| PE (28:1) * | C33H64NO8P | M+H+ | 634.4442 | 4.2 | Glycerophospholipids (GP) | Glycerophosphoethanolamines (PE) | Diacyl PE |
| PE (29:0)_5.4min | C34H68NO8P | M+H+ | 650.4755 | 5.35 | Glycerophospholipids (GP) | Glycerophosphoethanolamines (PE) | Diacyl PE |
| PE (29:0)_5.6min | C34H68NO8P | M+H+ | 650.4755 | 5.58 | Glycerophospholipids (GP) | Glycerophosphoethanolamines (PE) | Diacyl PE |
| PE (29:1)* | C34H66NO8P | M+H+ | 648.4599 | 4.5 | Glycerophospholipids (GP) | Glycerophosphoethanolamines (PE) | Diacyl PE |
| PE (30:0)_6.1min | C35H70NO8P | M+H+ | 664.4912 | 6.11 | Glycerophospholipids (GP) | Glycerophosphoethanolamines (PE) | Diacyl PE |
| PE (30:0)_6.4min | C35H70NO8P | M+H+ | 664.4912 | 6.41 | Glycerophospholipids (GP) | Glycerophosphoethanolamines (PE) | Diacyl PE |
| PE (30:0)_6.7min | C35H70NO8P | M+H+ | 664.4912 | 6.74 | Glycerophospholipids (GP) | Glycerophosphoethanolamines (PE) | Diacyl PE |
| PE (30:1)* | C35H68NO8P | M+H+ | 662.4755 | 5.4 | Glycerophospholipids (GP) | Glycerophosphoethanolamines (PE) | Diacyl PE |
| PE (31:0)_7.1min | C36H72NO8P | M+H+ | 678.5068 | 7.13 | Glycerophospholipids (GP) | Glycerophosphoethanolamines (PE) | Diacyl PE |

**Table S2.** Continued
| Lipid | Formula | Ion | m/z | t<sub>R</sub> | Lipid Category | Lipid Class | Lipid Subclass |
| --- | --- | --- | --- | --- | --- | --- | --- |
| PE (31:0)_7.4min | C36H72NO8P | M+H+ | 678.5068 | 7.40 | Glycerophospholipids (GP) | Glycerophosphoethanolamines (PE) | Diacyl PE |
| PE (31:1)* | C36H70NO8P | M+H+ | 676.4912 | 5.9 | Glycerophospholipids (GP) | Glycerophosphoethanolamines (PE) | Diacyl PE |
| PE (32:0)_8.2min | C37H74NO8P | M+H+ | 692.5229 | 8.18 | Glycerophospholipids (GP) | Glycerophosphoethanolamines (PE) | Diacyl PE |
| PE (32:0)_8.5min | C37H74NO8P | M+H+ | 692.5229 | 8.54 | Glycerophospholipids (GP) | Glycerophosphoethanolamines (PE) | Diacyl PE |
| PE (32:0)_9.0min | C37H74NO8P | M+H+ | 692.5229 | 8.96 | Glycerophospholipids (GP) | Glycerophosphoethanolamines (PE) | Diacyl PE |
| PE (32:1)_7.0min | C37H72NO8P | M+H+ | 690.5072 | 6.99 | Glycerophospholipids (GP) | Glycerophosphoethanolamines (PE) | Diacyl PE |
| PE (32:1)_7.2min | C37H72NO8P | M+H+ | 690.5072 | 7.22 | Glycerophospholipids (GP) | Glycerophosphoethanolamines (PE) | Diacyl PE |
| PE (33:0)_9.9min | C38H76NO8P | M+H+ | 706.5386 | 9.86 | Glycerophospholipids (GP) | Glycerophosphoethanolamines (PE) | Diacyl PE |
| PE (33:0)_10.0min | C38H76NO8P | M+H+ | 706.5386 | 10.03 | Glycerophospholipids (GP) | Glycerophosphoethanolamines (PE) | Diacyl PE |
| PE (33:1)_7.6min | C38H74NO8P | M+H+ | 704.5229 | 7.62 | Glycerophospholipids (GP) | Glycerophosphoethanolamines (PE) | Diacyl PE |
| PE (33:1)_8.1min | C38H74NO8P | M+H+ | 704.5229 | 8.05 | Glycerophospholipids (GP) | Glycerophosphoethanolamines (PE) | Diacyl PE |
| PE (34:1) | C39H76NO8P | M+H+ | 718.5386 | 9.23 | Glycerophospholipids (GP) | Glycerophosphoethanolamines (PE) | Diacyl PE |
| PE (P-25:0) | C30H60NO7P | M-H- | 576.4035 | 3.78 | Glycerophospholipids (GP) | Glycerophosphoethanolamines (PE) | 1-(1Z-alkenyl),2-acylPE (PE P) |
| PE (P-26:0)_4.1min | C31H62NO7P | M-H- | 590.4191 | 4.07 | Glycerophospholipids (GP) | Glycerophosphoethanolamines (PE) | 1-(1Z-alkenyl),2-acylPE (PE P) |
| PE (P-26:0)_4.3min | C31H62NO7P | M-H- | 590.4191 | 4.30 | Glycerophospholipids (GP) | Glycerophosphoethanolamines (PE) | 1-(1Z-alkenyl),2-acylPE (PE P) |
| PE (P-26:0)_4.4min | C31H62NO7P | M-H- | 590.4191 | 4.44 | Glycerophospholipids (GP) | Glycerophosphoethanolamines (PE) | 1-(1Z-alkenyl),2-acylPE (PE P) |
| PE (P-27:0)_4.7min | C32H64NO7P | M-H- | 604.4348 | 4.66 | Glycerophospholipids (GP) | Glycerophosphoethanolamines (PE) | 1-(1Z-alkenyl),2-acylPE (PE P) |
| PE (P-27:0)_4.9min | C32H64NO7P | M-H- | 604.4348 | 4.86 | Glycerophospholipids (GP) | Glycerophosphoethanolamines (PE) | 1-(1Z-alkenyl),2-acylPE (PE P) |
| PE (P-28:0)_5.3min | C33H66NO7P | M-H- | 618.4504 | 5.32 | Glycerophospholipids (GP) | Glycerophosphoethanolamines (PE) | 1-(1Z-alkenyl),2-acylPE (PE P) |
| PE (P-28:0)_5.6min | C33H66NO7P | M-H- | 618.4504 | 5.61 | Glycerophospholipids (GP) | Glycerophosphoethanolamines (PE) | 1-(1Z-alkenyl),2-acylPE (PE P) |
| PE (P-28:0)_5.8min | C33H66NO7P | M-H- | 618.4504 | 5.82 | Glycerophospholipids (GP) | Glycerophosphoethanolamines (PE) | 1-(1Z-alkenyl),2-acylPE (PE P) |

**Table S2.** Continued
| Lipid | Formula | Ion | m/z | t<sub>R</sub> | Lipid Category | Lipid Class | Lipid Subclass |
| --- | --- | --- | --- | --- | --- | --- | --- |
| PE (P-29:0)_6.2min | C34H68NO7P | M-H- | 632.4661 | 6.17 | Glycerophospholipids (GP) | Glycerophosphoethanolamines (PE) | 1-(1Z-alkenyl),2-acylPE (PE P) |
| PE (P-29:0)_6.4min | C34H68NO7P | M-H- | 632.4661 | 6.41 | Glycerophospholipids (GP) | Glycerophosphoethanolamines (PE) | 1-(1Z-alkenyl),2-acylPE (PE P) |
| PE (P-30:0)_7.0min | C35H70NO7P | M-H- | 646.4817 | 7.03 | Glycerophospholipids (GP) | Glycerophosphoethanolamines (PE) | 1-(1Z-alkenyl),2-acylPE (PE P) |
| PE (P-30:0)_7.4min | C35H70NO7P | M-H- | 646.4817 | 7.36 | Glycerophospholipids (GP) | Glycerophosphoethanolamines (PE) | 1-(1Z-alkenyl),2-acylPE (PE P) |
| PE (P-31:0) | C36H72NO7P | M-H- | 660.4974 | 8.50 | Glycerophospholipids (GP) | Glycerophosphoethanolamines (PE) | 1-(1Z-alkenyl),2-acylPE (PE P) |
| LPE (15:0) | C20H42NO7P | M-H- | 438.2627 | 1.14 | Glycerophospholipids (GP) | Glycerophosphoethanolamines (PE) | MonoacylPE (LPE) |
| LPE (16:0) | C21H44NO7P | M-H- | 452.2783 | 1.41 | Glycerophospholipids (GP) | Glycerophosphoethanolamines (PE) | MonoacylPE (LPE) |
| LPE (17:0) | C22H46NO7P | M-H- | 466.2940 | 1.68 | Glycerophospholipids (GP) | Glycerophosphoethanolamines (PE) | MonoacylPE (LPE) |
| LPE (P-13:0) | C18H38NO6P | M-H- | 394.2364 | 0.99 | Glycerophospholipids (GP) | Glycerophosphoethanolamines (PE) | 1Z-alkenylPE (LPE P) |
| LPE (P-14:0) | C19H40NO6P | M-H- | 408.2521 | 1.18 | Glycerophospholipids (GP) | Glycerophosphoethanolamines (PE) | 1Z-alkenylPE (LPE P) |
| LPE (P-15:0) | C20H42NO6P | M-H- | 422.2677 | 1.31 | Glycerophospholipids (GP) | Glycerophosphoethanolamines (PE) | 1Z-alkenylPE (LPE P) |
| LPE (P-16:0) | C21H44NO6P | M-H- | 436.2834 | 1.57 | Glycerophospholipids (GP) | Glycerophosphoethanolamines (PE) | 1Z-alkenylPE (LPE P) |
| PI (27:0) | C36H69O13P | M-H- | 739.4403 | 2.55 | Glycerophospholipids (GP) | Glycerophosphoinositols (PI) | diacyl PI |
| PI (28:0) | C37H71O13P | M-H- | 753.4560 | 2.82 | Glycerophospholipids (GP) | Glycerophosphoinositols (PI) | diacyl PI |
| PI (29:0) | C38H73O13P | M-H- | 767.4716 | 3.18 | Glycerophospholipids (GP) | Glycerophosphoinositols (PI) | diacyl PI |
| PI (30:0) | C39H75O13P | M-H- | 781.4873 | 3.42 | Glycerophospholipids (GP) | Glycerophosphoinositols (PI) | diacyl PI |
| PI (31:0) | C40H77O13P | M-H- | 795.5029 | 3.91 | Glycerophospholipids (GP) | Glycerophosphoinositols (PI) | diacyl PI |
| PS (28:0) | C34H66NO10P | M-H- | 678.4352 | 3.9 | Glycerophospholipids (GP) | Glycerophosphoserines (PS) | diacyl PS |
| PS (29:0) | C35H68NO10P | M-H- | 692.4508 | 4.6 | Glycerophospholipids (GP) | Glycerophosphoserines (PS) | diacyl PS |
| PS (30:0) | C36H70NO10P | M-H- | 706.4665 | 5.1 | Glycerophospholipids (GP) | Glycerophosphoserines (PS) | diacyl PS |
| PS (31:0) | C37H72NO10P | M-H- | 720.4821 | 5.3 | Glycerophospholipids (GP) | Glycerophosphoserines (PS) | diacyl PS |

**Table S2.** Continued
| Lipid | Formula | Ion | m/z | t<sub>R</sub> | Lipid Category | Lipid Class | Lipid Subclass |
| --- | --- | --- | --- | --- | --- | --- | --- |
| sphinganine C15 | C15H33NO2 | M+H+ | 260.2584 | 1.04 | Sphingolipids (SP) | Sphingoid bases | Sphinganines |
| sphinganine C16 | C16H35NO2 | M+H+ | 274.2742 | 1.24 | Sphingolipids (SP) | Sphingoid bases | Sphinganines |
| sphinganine C17 | C17H37NO2 | M+H+ | 288.2899 | 1.34 | Sphingolipids (SP) | Sphingoid bases | Sphinganines |
| sphinganine C18 | C18H39NO2 | M+H+ | 302.3056 | 1.67 | Sphingolipids (SP) | Sphingoid bases | Sphinganines |
| sphinganine C19 | C19H41NO2 | M+H+ | 316.3212 | 1.87 | Sphingolipids (SP) | Sphingoid bases | Sphinganines |
| sphinganine C20 | C20H43NO2 | M+H+ | 330.3369 | 2.33 | Sphingolipids (SP) | Sphingoid bases | Sphinganines |
| hydroxy sphinganine C18 | C18H39NO3 | M+H+ | 318.3003 | 1.31 | Sphingolipids (SP) | Sphingoid bases | Sphinganines |
| hydroxy sphinganine C19 | C19H41NO3 | M+H+ | 332.3159 | 1.45 | Sphingolipids (SP) | Sphingoid bases | Sphinganines |
| keto-sphinganine C15 | C15H31NO2 | M+H+ | 258.2428 | 1.04 | Sphingolipids (SP) | Sphingoid bases | Sphinganines |
| keto-sphinganine C16 | C16H33NO2 | M+H+ | 272.2584 | 1.21 | Sphingolipids (SP) | Sphingoid bases | Sphinganines |
| keto-sphinganine C17 | C17H35NO2 | M+H+ | 286.2741 | 1.31 | Sphingolipids (SP) | Sphingoid bases | Sphinganines |
| keto-sphinganine C18 | C18H37NO2 | M+H+ | 300.2897 | 1.60 | Sphingolipids (SP) | Sphingoid bases | Sphinganines |
| keto-sphinganine C19 | C19H39NO2 | M+H+ | 314.3054 | 1.80 | Sphingolipids (SP) | Sphingoid bases | Sphinganines |
| keto-sphinganine C20 | C20H41NO2 | M+H+ | 328.3210 | 2.26 | Sphingolipids (SP) | Sphingoid bases | Sphinganines |
| hydroxy keto-sphinganine C18 | C18H37NO2 | M+H+ | 316.2846 | 1.25 | Sphingolipids (SP) | Sphingoid bases | Sphinganines |
| hydroxy keto-sphinganine C19 | C19H39NO2 | M+H+ | 330.3003 | 1.41 | Sphingolipids (SP) | Sphingoid bases | Sphinganines |
| deoxy-sphinganine C17 | C17H37NO | M+H+ | 272.2948 | 1.48 | Sphingolipids (SP) | Sphingoid bases | Sphingoid base analogs |
| deoxy-sphinganine C18 | C18H39NO | M+H+ | 286.3104 | 1.84 | Sphingolipids (SP) | Sphingoid bases | Sphingoid base analogs |
| deoxy-sphinganine C19 | C19H41NO | M+H+ | 300.3261 | 2.09 | Sphingolipids (SP) | Sphingoid bases | Sphingoid base analogs |
| oxidised dihydroCer C34 | C34H67NO4 | M+Na | 576.4968 | 6.80 | Sphingolipids (SP) | Ceramides (Cer) | Dihydroceramides (dihydroCer) |
| dihydroCer C32_5.0min | C32H65NO4 | M+Na | 550.4815 | 5.02 | Sphingolipids (SP) | Ceramides (Cer) | Dihydroceramides (dihydroCer) |
| dihydroCer C32_5.2min | C32H65NO4 | M+Na | 550.4815 | 5.19 | Sphingolipids (SP) | Ceramides (Cer) | Dihydroceramides (dihydroCer) |
| dihydroCer C32_5.4min | C32H65NO4 | M+Na | 550.4815 | 5.39 | Sphingolipids (SP) | Ceramides (Cer) | Dihydroceramides (dihydroCer) |
| dihydroCer C33_5.8min | C33H67NO4 | M+H+ | 542.5146 | 5.77 | Sphingolipids (SP) | Ceramides (Cer) | Dihydroceramides (dihydroCer) |

**Table S2.** Continued
| Lipid | Formula | Ion | m/z | t<sub>R</sub> | Lipid Category | Lipid Class | Lipid Subclass |
| --- | --- | --- | --- | --- | --- | --- | --- |
| dihydroCer C33_6.0min | C33H67NO4 | M+H+ | 542.5146 | 5.97 | Sphingolipids (SP) | Ceramides (Cer) | Dihydroceramides (dihydroCer) |
| dihydroCer C33_6.2min | C33H67NO4 | M+H+ | 542.5146 | 6.21 | Sphingolipids (SP) | Ceramides (Cer) | Dihydroceramides (dihydroCer) |
| dihydroCer C34_6.6min | C34H69NO4 | M+H+ | 556.5303 | 6.63 | Sphingolipids (SP) | Ceramides (Cer) | Dihydroceramides (dihydroCer) |
| dihydroCer C34_6.9min | C34H69NO4 | M+H+ | 556.5303 | 6.90 | Sphingolipids (SP) | Ceramides (Cer) | Dihydroceramides (dihydroCer) |
| dihydroCer C34_7.2min | C34H69NO4 | M+H+ | 556.5303 | 7.19 | Sphingolipids (SP) | Ceramides (Cer) | Dihydroceramides (dihydroCer) |
| dihydroCer C35_7.7min | C35H71NO4 | M+H+ | 570.5459 | 7.66 | Sphingolipids (SP) | Ceramides (Cer) | Dihydroceramides (dihydroCer) |
| dihydroCer C35_8.0min | C35H71NO4 | M+H+ | 570.5459 | 8.01 | Sphingolipids (SP) | Ceramides (Cer) | Dihydroceramides (dihydroCer) |
| dihydroCer C36 | C36H73NO4 | M+H+ | 584.5616 | 8.84 | Sphingolipids (SP) | Ceramides (Cer) | Dihydroceramides (dihydroCer) |
| deoxy-dihydroCer C34_7.5min | C34H69NO3 | [M+Na] | 562.5179 | 7.48 | Sphingolipids (SP) | Ceramides (Cer) | Dihydroceramides (dihydroCer) |
| deoxy-dihydroCer C34_7.8min | C34H69NO3 | [M+Na] | 562.5179 | 7.76 | Sphingolipids (SP) | Ceramides (Cer) | Dihydroceramides (dihydroCer) |
| deoxy-dihydroCer C35_8.7min | C35H71NO3 | [M+Na] | 576.5332 | 8.67 | Sphingolipids (SP) | Ceramides (Cer) | Dihydroceramides (dihydroCer) |
| deoxy-dihydroCer C35_9.0min | C35H71NO3 | [M+Na] | 576.5332 | 9.03 | Sphingolipids (SP) | Ceramides (Cer) | Dihydroceramides (dihydroCer) |
| deoxy-dihydroCer C36 | C36H73NO3 | [M+Na] | 590.5492 | 9.98 | Sphingolipids (SP) | Ceramides (Cer) | Dihydroceramides (dihydroCer) |
| Cer PE C33 | C35H73N2O7P | M+H+ | 665.5232 | 4.93 | Sphingolipids (SP) | Phosphosphingolipids | dihydroCer PE (Cer PE) |
| Cer PE C34_5.3min | C36H75N2O7P | M+H+ | 679.5389 | 5.26 | Sphingolipids (SP) | Phosphosphingolipids | dihydroCer PE (Cer PE) |
| Cer PE C34_5.5min | C36H75N2O7P | M+H+ | 679.5389 | 5.48 | Sphingolipids (SP) | Phosphosphingolipids | dihydroCer PE (Cer PE) |
| Cer PE C34_5.7min | C36H75N2O7P | M+H+ | 679.5389 | 5.68 | Sphingolipids (SP) | Phosphosphingolipids | dihydroCer PE (Cer PE) |
| Cer PE C35_6.0min | C37H77N2O7P | M+H+ | 693.5545 | 6.04 | Sphingolipids (SP) | Phosphosphingolipids | dihydroCer PE (Cer PE) |
| Cer PE C35_6.3min | C37H77N2O7P | M+H+ | 693.5545 | 6.31 | Sphingolipids (SP) | Phosphosphingolipids | dihydroCer PE (Cer PE) |
| Cer PE C36 | C38H79N2O7P | M+H+ | 707.5702 | 6.99 | Sphingolipids (SP) | Phosphosphingolipids | dihydroCer PE (Cer PE) |
| Cer PI C33 | C39H78NO12P | M+H+ | 784.5334 | 2.93 | Sphingolipids (SP) | Phosphosphingolipids | dihydroCer PI (Cer PI) |

**Table S2.** Continued
| Lipid | Formula | Ion | m/z | t<sub>R</sub> | Lipid Category | Lipid Class | Lipid Subclass |
| --- | --- | --- | --- | --- | --- | --- | --- |
| Cer PI C34 | C40H80NO12P | M+H+ | 798.5495 | 3.11 | Sphingolipids (SP) | Phosphosphingolipids | dihydroCer PI (Cer PI) |
| Cer PI C35 | C41H82NO12P | M+H+ | 812.5652 | 3.57 | Sphingolipids (SP) | Phosphosphingolipids | dihydroCer PI (Cer PI) |
| Cer PI C36 | C42H84NO12P | M+H+ | 826.5809 | 3.84 | Sphingolipids (SP) | Phosphosphingolipids | dihydroCer PI (Cer PI) |
| α-Gal Cer C33 | C39H77NO9 | M+H+ | 704.5671 | 4.99 | Sphingolipids (SP) | Neutral glycosphingolipids | Simple Glc series |
| α-Gal Cer C34_5.5min | C40H79NO9 | M+H+ | 718.5828 | 5.51 | Sphingolipids (SP) | Neutral glycosphingolipids | Simple Glc series |
| α-Gal Cer C34_5.7min | C40H79NO9 | M+H+ | 718.5828 | 5.70 | Sphingolipids (SP) | Neutral glycosphingolipids | Simple Glc series |
| α-Gal Cer C35_6.3min | C41H81NO9 | M+H+ | 732.5984 | 6.34 | Sphingolipids (SP) | Neutral glycosphingolipids | Simple Glc series |
| α-Gal Cer C35_6.6min | C41H81NO9 | M+H+ | 732.5984 | 6.60 | Sphingolipids (SP) | Neutral glycosphingolipids | Simple Glc series |
| α-Gal Cer C36 | C42H83NO9 | M+H+ | 746.6141 | 7.33 | Sphingolipids (SP) | Neutral glycosphingolipids | Simple Glc series |
| 16:0 SM | C39H79N2O6P | M+HCOO- | 747.5658 | 6.17 | Sphingolipids (SP) | Internal Standard | Internal Standard |
| 16:0 SM | C39H79N2O6P | M+H+ | 703.5749 | 6.17 | Sphingolipids (SP) | Internal Standard | Internal Standard |

DG: diacylglycerols; C: carbon; FL: flavolipin; mono: monoacyl; GlyL: glycine lipid; Gal: galactosyl; SM: sphingomyelin
\* May contain more than one peak/isomer
Where possible LipidMaps classification & nomenclature is used throughout.
GP abbreviation is based on sum composition which accounts for the total number of carbons and double bonds in the fatty acyl chains (number of carbons: number double bonds). For example, PE (30:0) denotes a diacyl PE whereby the sum of carbons in the 2 acyl chains is 30 and both acyl chains are saturated (no double bond); possibilities include a C17:0 and C13:0; C15:0 and C15:0; C14:0 and C16:0 and so on. Fatty acyl chains may also be branched/unbranched. Where there are more than one chromatographic peak GP are also distinguished on their elution time (min), for example PE (30:0)\_6.1 min, and indicates the possibilities of numerous combinations of fatty acyls (chain lengths and branching)
*Bacteroides* SP abbreviation is based on the combined total number of carbons on the sphinganine backbone and the fatty acyl chain. Where there is more than one peak, they are also distinguished based on their elution or retention time (t<sub>R</sub> in min). For example, Cer PE C34\_5.3min has a combined total of 34 carbons in the sphinganine backbone and fatty acyl chain and elutes at 5.3 min. Like GP there are numerous combinations of sphinganine and fatty acyl chain length as well as branching and non-branching on the sphinganine and/or the fatty acyl component. There is evidence to suggest that debranched SP elute first with subsequent elution of mono-branched and unbranched isomers (Oh et al. 2021).

